# Designed Minibinders Rewire Receptor Signaling to Enable Functional Human Myogenic Reprogramming

**DOI:** 10.64898/2026.04.26.720818

**Authors:** Riya Keshri, Aidan Jessop, Alexander Robinson, Zachary Foreman, Philip Barrett, Gabriela Reyes, Aditya KrishnaKumar, Saurav Kumar, Ashish Phal, Ethan Narog, Melodie Chiu, Shruti Jain, Xinru Wang, David Lee, Marc Exposit, Mohamad Adedi, Alec ST Smith, Sanjay Srivatsan, Jay Shendure, Julie Mathieu, David L Mack, David Baker, Hannele Ruohola-Baker

## Abstract

Sarcopenia, loss of muscle mass is a considerable health burden that demands immediate societal attention and calls for new regenerative strategies. Direct myogenic somatic cell reprogramming, a potential muscle regeneration method is constrained by an inability to control the signaling logic that governs cell fate. Here, we show that this barrier can be overcome using AI-designed receptor modulators. Screening de novo minibinders, we identify a synthetic protein cocktail, C6-DPC, that drives efficient human fibroblast-to-muscle transdifferentiation with robust structural and metabolic maturation. C6-DPC reprograms extracellular signaling by activating pro-myogenic FGFR1/2-c pathways while suppressing anti-myogenic inputs through ALK1 and TGFBR2; targeted depletion of ALK1 is sufficient to lower the reprogramming barrier. Inflammatory signaling via gp130 emerges as a dominant checkpoint, and its inhibition further enhances fibroblast-to-muscle conversion. C6-DPC treated engineered tissues generate high twitch and tetanic forces in both wild-type and dystrophin-deficient human cells. These findings demonstrate programmable control of receptor signaling as a strategy to direct myogenic cell fate and generate functional human muscle, providing a foundation for regenerative approaches to muscle loss and disease.

## Introduction

Skeletal muscle strength and function begin to decline as early as the third decade of life, initiating a progressive trajectory toward sarcopenia, a major and growing health burden in the aging population (Pabla et al., 2024). Under physiological conditions, muscle homeostasis is sustained by satellite cells; however, their regenerative capacity diminishes sharply with age and can be further compromised by severe trauma, surgical injury, and genetic disorders such as duchenne muscular dystrophy (DMD). Moreover, the clinical success resulting in population level use of GLP-1 receptor agonists (e.g., semaglutide, tirzepatide) can involve a significant reduction in lean body mass (LBM), which may exacerbate or trigger early sarcopenia (Scott et al.,2025; Langer et al., 2026).

Direct cellular reprogramming from one adult cell type to another has shown to occur in nature (Wu et al., 2012; Thorel et al., 2010). This process has recently been harnessed for regenerative medicine, offering a path to bypass the complexities of pluripotency while providing a source of patient specific functional cells. The conversion of fibroblasts into myogenic lineages via the ectopic expression of MyoD was the seminal demonstration of cell fate plasticity (Weintraub et al., 1987; Morris & Daley, 2013). However, despite decades of research, the therapeutic application of direct reprogramming is still hampered by low conversion efficiency (Cacchiarelli et al., 2015; Kim et al., 2022). Although MyoD established the principle of cell fate plasticity, its practical application remains limited by inefficient conversion and incomplete maturation. These limitations reflect a fundamental issue: cell fate is not governed by transcription factors alone, but by a complex extracellular signaling environment that needs to be controlled (Boularaoui et al., 2018; Cacchiarelli et al., 2018). Competing cues, including pro-fibrotic pathways actively oppose myogenic conversion, while current small-molecule approaches lack the specificity to resolve these conflicts. Furthermore, restricted chromatin accessibility limits MyoD binding to key regulatory elements and results in incomplete transcriptional activation of the myogenic program (Manandhar et al., 2017; Kolundzic et al., 2018; Liu et al., 2018). With small molecule approaches, dedifferentiation into induced myogenic progenitor cells (iMPCs) has also been explored, however these cells can remain highly proliferative (Yagi et al., 2021; Kim et al., 2022).

Recent breakthroughs in AI based molecular modeling, such as RFDiffusion and BindCraft, have introduced a new class of therapeutic agents: synthetic minibinders (Watson et al., 2023; Pacesa et al., 2025). These *de novo* designed proteins can be engineered with exquisite specificity down to the receptor isoform level and can function as either potent antagonists in monomeric form or "superagonists" when organized into high valency oligomeric scaffolds (Cao et al, 2022; Edman et al., 2024; Zhao et al., 2021). Unlike natural ligands, these minibinders can be displayed on rigid or flexible scaffolds to precisely tune signaling geometries. Recent work has already demonstrated their power; for example, a designed FGFR1/2-c agonist can switch the fate of pluripotent stem cells between endothelial and pericyte states (Edman et al. 2024).

In this study, we screened a library of AI-designed proteins to identify novel designed protein cocktails that can overcome the efficiency bottleneck of myogenic conversion. We report that a strategic combination of minibinders specifically targeting the activation of FGFR1/2-c alongside the inhibition of TGFBRII and ALK1, significantly increases the efficiency and maturation of human fibroblast derived muscle cells. In parallel, inhibition of inflammatory signaling via gp130 removes a dominant barrier to conversion identifying inflammation as a dominant checkpoint in reprogramming. Our results demonstrate that these synthetic protein cocktails not only refine the reprogramming process but also facilitate the functional rescue of DMD muscle force phenotypes, marking a new era in precision regenerative therapy.

## Results

### Designed receptor modulatory protein cocktail (DPC) enhances the skeletal muscle direct reprogramming

In this study we utilize designed proteins to target receptor tyrosine kinases or receptor serine/threonine kinases to make the human myogenic cell fate conversion efficient. We engineered neonatal skin fibroblasts (HFFs, Human Foreskin Fibroblasts) to generate an inducible MyoD line (HFF-iMYOD) under the control of a Tet-on promoter **(Figs. EV1A-B)**. The continuous doxycycline-induced overexpression of MyoD in HFF-iMYOD for 7 and 14 days resulted in a low conversion of fibroblasts to the myogenic lineage (**Figs. EV1C-G)**. Transcriptomic analysis revealed three distinct cell populations; resting fibroblasts maintaining baseline markers (MME, CXCL12), a population of activated myofibroblasts enriched for fibrotic markers (FN1, IGFBP5), and a reprogrammed cluster expressing muscle structural and functional genes (ACTC1, CHRNA1) **(Fig. EV1H)**.

We analyzed a set of barrier genes (ID1, ID2, ID3, TGFB1, TGFB2, TGFBR1, TWIST1, TWIST2, ZEB1, SNAI1, SNAI2, HES1, HEY1, NOTCH1, WNT5A) previously reported to inhibit myogenic differentiation through repression of MyoD activity, activation of Epithelial-Mesenchymal Transition (EMT) programs, or maintenance of progenitor states via TGF-β and Notch signaling pathways (Benezra et al., 1990; Spicer et al., 1996; Buas et al., 2009; Soleimani et al., 2012; von Maltzahn et al., 2012). Our analysis identified a hierarchical "brake" system categorized into four distinct regulatory drivers that actively antagonize myogenic transdifferentiation **(Fig. EV1I)**. The most potent barrier was the structural reinforcement of the extracellular matrix (ECM), characterized by strong positive correlations with *COL1A2* , *SPARC*, and *COL3A1* suggesting the physical rigidity and integrin signaling of the matrix structurally anchor cells to a non-myogenic identity. Activated myofibroblast markers, including *TAGLN*, *CTGF*, *INHBA*, and *POSTN*, showed significant positive correlation with the barrier score, confirming that a competing pro-fibrotic program directly diverts cells from the skeletal muscle fate **(Fig. EV1I)**. We identified active molecular inhibitors of myogenesis, specifically *IGFBP5*, *IGFBP7*, and *ID3* **(Fig. EV1I)**, and also identified a novel category of myogenic inhibitors, regulators of cellular stress and inflammation (*IFI6*, *STAT3*, and *NFKB1*). Collectively, these results map a multi-layered roadblock to myogenesis, where successful reprogramming requires the simultaneous suppression of structural ECM traps, competing fibrotic lineages, transcriptional repressors, and inflammatory stress states **(Fig. EV1I)**.

To date, numerous biological ligands including Follistatin, GDF8, FGF2, GDF11, GDF15, hGH, TMSB4X, BMP4, BMP7, IL-6, and TNF-a have been evaluated for their ability to enhance MyoD induced transdifferentiation process, but none have proven successful (Abdel-Raouf et al., 2021). However, these factors predominantly promote cellular proliferation or function as negative regulators that further impede myogenic conversion. To identify novel potent signaling pathways beneficial for direct myoblast reprogramming, we established a streamlined, defined serum-free screening assay and screened for novel AI-designed proteins that enhanced the direct reprogramming. We conducted a temporal screen to define the minimal Dox exposure required to prime the fibroblasts for myogenic reprogramming. Following induction, cells were transitioned into serum-free media for various periods to observe the progression of differentiation **(Fig. EV1J,K)**. Our time-course analysis revealed that a 4-day Dox pulse (D4) followed by a 4-day release into serum-free media (D8) was sufficient to drive consistent, albeit still low myogenic conversion **(Fig.1A)**.

**Figure 1.**
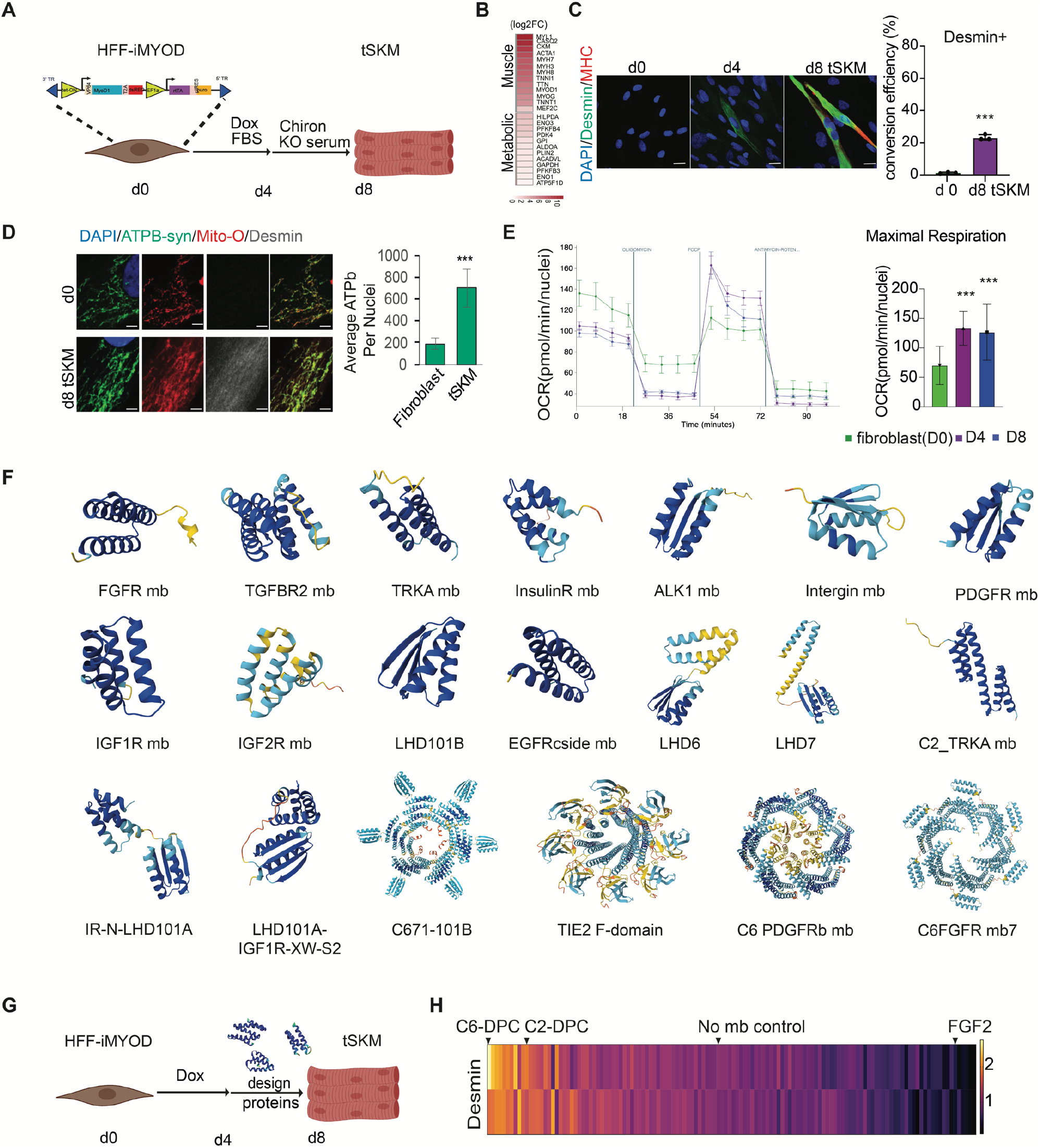
Direct reprogramming of human foreskin fibroblast into skeletal muscles. (A) Schematic showing Tet-on inducible MyoD Human Foreskin FIbroblast were treated with Doxycycline for 4 days, then treated with designed proteins & chiron in KO serum media for the following 4 days to reprogram into muscle cells, where immunofluorescent staining for muscle markers was performed. (B) Heatmap showing 8 days tSKM versus fibroblast bulk sequencing data for muscle markers and metabolic gene expression. (C) Day 8 (D4>8) reprogrammed tSKM shows muscle markers staining for Desmin (green), Myosin heavy chain (red), and DAPI (blue). A bar graph for conversion efficiency (desmin+) of uninduced HFFs vs. day 8 tSKM. Scale bar, 30um (n=3). (D) Confocal images of uninduced fibroblast and day 8 tSKM stained with mitochondrial marker ATP-b synthase (green), mito-Orange (red) uptake, Desmin (grey) and DAPI (blue). Graph shows quantification of ATP-b synthase intensity in uninduced fibroblast, day 8 no mb tSKM, day 8 DPC treated tSKM. Scale bar, 5um (n=3). (E) Normalized OCR measure in uninduced fibroblast, day 4 and day 8 tSKM. Maximum respiration measured in uninduced fibroblast, day 4 and day 8 tSKM (n=3). (F) Few examples of Alphafold3 predicted structure of design binders used in the muscle transdifferentiation. Antagonists: FGFR mb, TGFBR2 mb, TRKA mb, InsulinR mb, ALK1 mb, Integrin mb, PDGFR mb, IGF1R mb, IGF2R mb, LHD101B, and EGFRcside mb. Agonists: C2_TRKA mb, IR-N-LHD101A, C671-101B, TIE2 F domain mb, C6-PDGFRb mb, and C6FGFR mb7. Scaffolds: LHD6 and LHD7 (G) schematic showing minibinder screen in skeletal muscle transdifferentiation. (H) A heatmap showing two independent screens of minibinders in day 8 tSKM. Where desmin intensity in two independent primary screens of designed proteins at 100 nM concentration in iMYODHFF has been plotted. Data are presented as mean ± SD with two-tailed unpaired Student’s *t*-tests. Where, *ns* for not significant \**p* < 0.05; \*\**p* < 0.01; \*\*\**p* < 0.001 and \*\*\*\**p*<0.0001.

**Figure 2.**
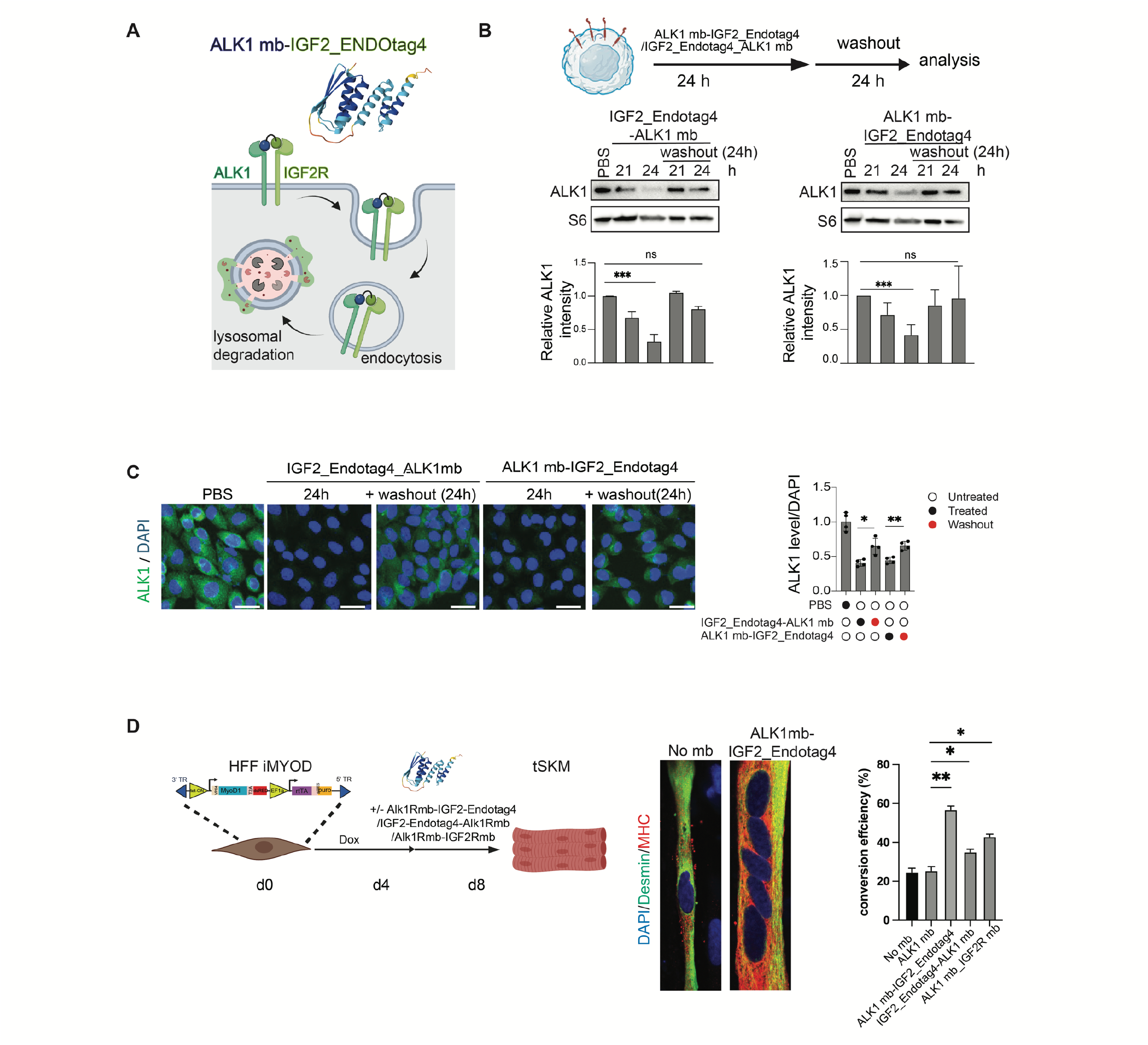
Strong degradation of ALK1R augments transdifferentiation. (A) Schematic of ALK1 mb-IGF2R_Endotag4 fusion directed endosomal internalization and lysosome mediated degradation of ALK1 receptor together with IGF2R. (B) Immunoblots and bar graphs of CHO cells overexpressing the human ALK1 are treated with the ALK1 mb-IGF2R_Endotag4mb or IGF2R_Endotag4mb-ALK1 mb for 24 hours and washed out for another 24 hours. The bar graph shows levels of ALK1 across multiple replicates (n=3). (C) Immunofluorescence images of ALK1 (green), phalloidin (red) and DAPI (blue) staining in the CHO cells exogenously expressing human ALK1. Scale bar, 50um (n=4 ). (D) Immunofluorescence images of reprogrammed tSKM upon no mb (control) or ALK1-IGF2R_Endotag4 treatment and schematically stained for Desmin (green), Myosin heavy chain (red), DAPI (blue). A bar graph showing fibroblast to muscle conversion efficiency is significantly upregulated in ALK1-IGF2R_Endotag4 treatment. Scale bar, 15um (n=3). Data are presented as mean ± SD with two-tailed unpaired Student’s *t*-tests. Where, *ns* for not significant \**p* < 0.05; \*\**p* < 0.01; \*\*\**p* < 0.001 and \*\*\*\**p*<0.0001.

Under these stable D8 conditions, approximately 20% of the population expressed the intermediate filament Desmin, Myosin heavy chain, Myogenin, alpha-actinin **(Figs. 1B-C, and EV1L)**. Additionally, we observed significant metabolic remodeling, a hallmark of the transition from a fibroblastic state to a mature myogenic oxidative profile **(Figs. 1D-E)**. The Desmin⁺ cells showed elevated ATPB expression (**Fig. 1D**) and both D4 and D8 stages exhibited significantly enhanced mitochondrial maximal respiration compared to the starting fibroblast population (**Fig. 1E**). Thus, the D8 state captures both myogenic markers and the essential metabolic switch for functional muscle. However, conversion efficiency remained low (∼20%, **Fig. 1C**), making this an ideal platform to screen AI-designed minibinders with improved myogenic capacity.

Leveraging recent advances in computational protein design, we hypothesized that precise outside-in signaling control using engineered minibinders and agonists could overcome fate stabilizing barriers and promote productive myogenic reprogramming (**Fig. 1F) (**Keshri et al., 2024). We therefore employed an optimized D8 reprogramming paradigm, in which minibinders and designed agonists were applied under serum-free conditions between day 4 and day 8, a critical window for fate stabilization (**Figs. 1G-H)**.

We performed focused screens of 130 designed minibinder(mb) and mb-scaffold combinations (for example FGFR, ALK1, PDGFR, InsulinR, IGF1R, TGFBRII, and EGFR mb as monomeric antagonists and engineered agonists including FGFR-C6, TRKA-C2, Tie2-H8, and EGFR) (**Fig. 1F, Table EV1)** (Cao et al. 2021; Edman et al.,2024; Schlichthaerle et al., 2025; Zhao et al., 2021). We analyzed the desmin intensity of each condition as a readout for myogenic reprogramming **(Fig. 1H, Table EV2)**. From this screen, we identified two key design protein cocktails (DPC) that significantly enhanced myogenic conversion: (I) C6-DPC (TGFBRIImb antagonist + ALK1mb antagonist + FGFR1/2c-C6 agonist) and (II) C2-DPC (TGFBRIImb antagonist + ALK1mb antagonist + TRKA-C2 agonist) where ALK1 and TGFBR2 pathway inhibition remains central in both the DPCs **(Fig. 3A)**.

**Figure 3.**
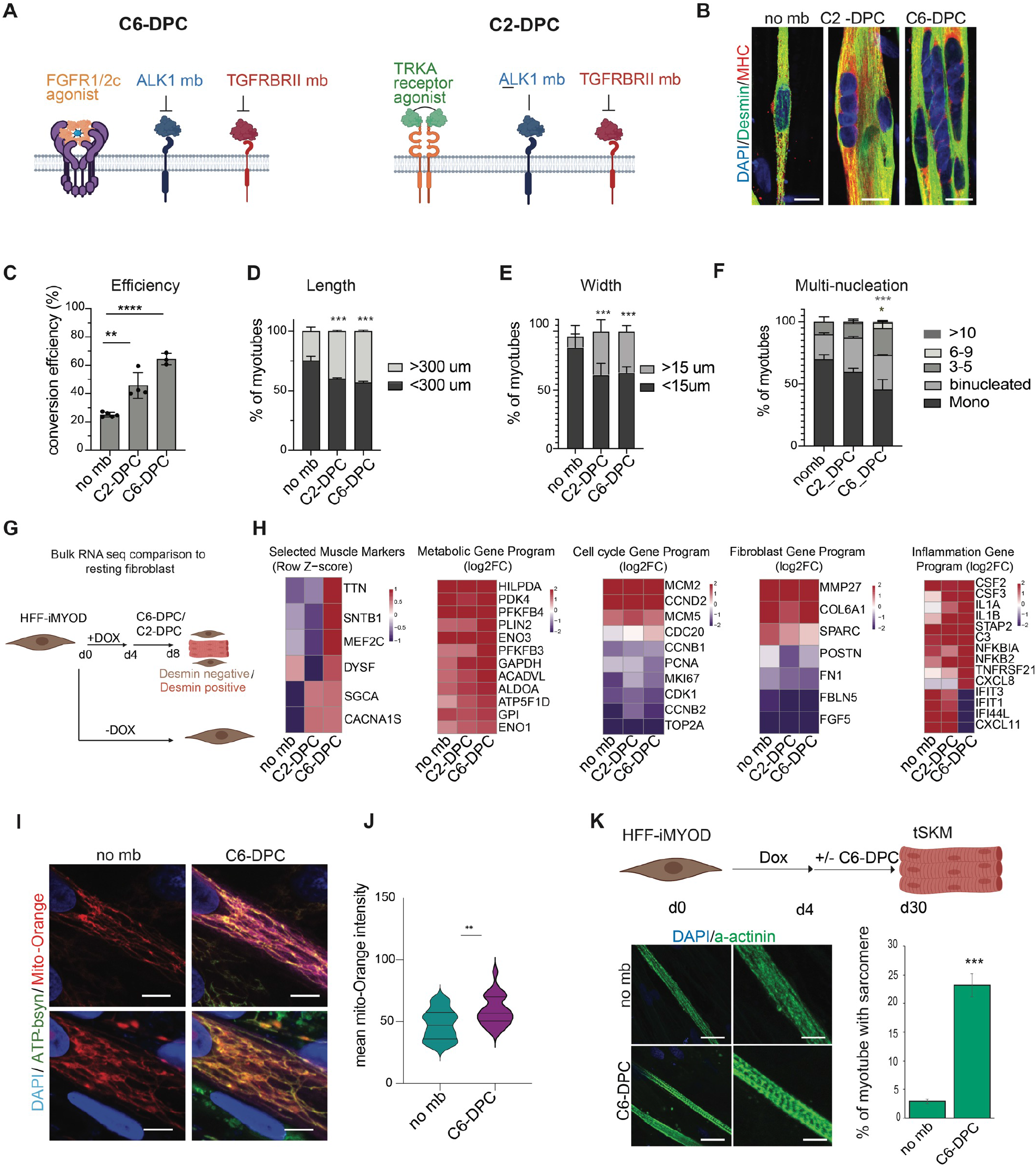
Designed protein cocktail enhanced myogenic transdifferentiation. (A) Schematic showing the two key designed protein cocktails, C6-DPC and C2-DPC. (B) Immunofluorescence images showing no mb, C2-DPC, and C6-DPC treated day 8 old tSKM stained for Desmin (green), MHC (red) and DAPI (blue). Scale bar,15um. (C) Bar graph showing conversion efficiency in C2-DPC and C6-DPC treated fibroblasts (n=3 ). (D) Length of myotubes in C2-DPC and C6-DPC treated tSKM (n=3 ). (E) Width of myotubes in C2-DPC and C6-DPC treated tSKM (n=3 ). (F) Percentage multinucleation of myotubes in C6-DPC treated tSKM (n=3 ). (G) Schematic showing Bulk RNA sequencing of tSKM and the mononucleated unconverted cells. (H) Heatmap showing muscle, metabolic, Cell cycle gene, fibroblast and inflammation markers expression at transcript level in no mb (control), C2-DPC, C6-DPC treatments versus fibroblast at day 8 conversion (RNA-seq data: GEO GSE330553). (I) Confocal images of uninduced fibroblast, day 8 no mb tSKM, day 8 C6-DPC treated tSKM stained for mitochondrial marker ATP-B synthase (green), mito-Orange (red) uptake, Desmin (magenta) and DAPI (blue). scale bar,5um. (J) Violin plot showing mito-Orange intensity in day 8 no mb tSKM, day 8 C6-DPC treated tSKM (n=_3). (K) Schematic of 30 days myotube maturation experiment. Sarcomere formation analysed with alpha-actinin staining (green) in day 30 no mb tSKM and C6-DPC treated tSKM. Scale bar,15um and 5um (n=3). Data are presented as mean ± SD with two-tailed unpaired Student’s *t*-tests. Where, *ns* for not significant \**p* < 0.05; \*\**p* < 0.01; \*\*\**p* < 0.001 and \*\*\*\**p*<0.0001.

### ALK1 signaling acts as barrier to efficient myogenic transdifferentiation

To validate the mechanism of the designed receptor modulatory cocktails, we focused on the inhibition of the Alk1 pathway. To test ALK1 inhibition by an alternative modality we utilized the IGF2_EndoTag4 minibinder which facilitates the endosomal internalization and subsequent lysosomal degradation of bound ligand complexes **(Fig. 2A**; Huang et. al., 2025). We therefore engineered an IGF2_EndoTag4-ALK1mb fusion designed to deplete ALK1 protein from the plasma membrane. We tested two orientations of the fusion proteins ALK1mb: IGF2_EndoTag4 mb (A1IE) and IGF2_EndoTag4 mb: ALK1mb (IEA1). To assess their efficacy, we treated Chinese Hamster Ovary (CHO) - human ALK1 cells for 24 hours and observed that both A1IE and IEA1 fusions induced a significant reduction in hALK1 protein levels **(Figs. 2B-C)**. ALK1 levels returned close to baseline after 24 hours removal of the fusions, confirming the specificity and reversibility of the EndoTag4 system **(Fig. 2B-C)**.

We tested the A1IE in the transdifferentiation assay **(Fig. 2D)**. Importantly, A1IE treatment significantly increased myogenic transdifferentiation efficiency, further supporting the conclusion that repression of Alk1 signaling pathway enhances myogenic transdifferentiation **(Fig. 2D)**,

The higher activity of ALK1mb–IGF2-EndoTag4 compared with ALK1mb alone indicates that depletion of ALK1 from the cell surface lowers the reprogramming barrier more effectively than antagonizing its activity by a minibinder, which we further confirmed through ALK1 shRNA knockdown **(Figs. EV1M)**. More broadly, our results establish AI-designed minibinder-driven targeted protein degradation as a powerful strategy to remove inhibitory signaling nodes. Using this approach, we identify ALK1 and TGFBRII as major molecular roadblocks to efficient muscle transdifferentiation and demonstrate that their programmable elimination can unlock latent myogenic potential.

### C6-DPC and C2-DPC accelerate skeletal muscle maturation

We analyzed the two powerful minibinder cocktails, C2-DPC and C6-DPC **(Fig. 3A).** Desmin and myosin heavy-chain staining showed more extensive myotube formation with both cocktails than in control cultures **(Fig. 3B)**. Comparison of individual components and their combinations using desmin-positive myotube area coverage and myogenic conversion efficiency identified C6-DPC as the most effective cocktail among the conditions tested **(Figs. 3C, EV2A-C).** These results identify combined FGFR1/2c stimulation and TGFBRII/ALK1 antagonism (C6-DPC) as a promising strategy for enhancing MYOD-driven myogenic conversion. C6-DPC treatment increased fibroblast conversion to the myogenic lineage over three-fold compared to the ∼20% baseline observed in standard conditions **(Fig. 3C)**. There was also a significant increase in both the length and width of the resulting myotubes, indicating more robust protein synthesis and sarcomeric assembly **(Figs. 3D-E, EV2D)**. Both C2-DPC and C6-DPC treated cells showed a significant increase in fusion index, characterized by the presence of multiple nuclei within a single continuous cytoplasm, a critical step for functional muscle contraction **(Figs. 3F, and EV2E)**. While both cocktails outperformed the control, the C6-DPC (FGFR1/2-c agonist + ALK1 and TGFBRII antagonists) consistently produced the most mature phenotypes.

### C6-DPC enhances myogenic remodeling

Transcriptomic analysis **(Fig. 3G)** revealed that both C2-DPC and C6-DPC induce MyoD-mediated fibroblast reprogramming leading to muscle marker expression (MYH7, ACTA1, TTN, MYL4, CASQ2 etc) and reduced expression of fibroblast markers (FBLN5, FN1, FGF5) **(Figs. 3H and EV3A, Table EV3)**. Both C2-DPC and C6-DPC conditions further induced a myogenic reprogramming response, as demonstrated by the upregulation of skeletal muscle associated genes (LMOD3, MYL6B, COX6A2, PDLIM4) involved in sarcomere assembly and contractile function **(Fig. EV3B).** Furthermore, metabolic genes were upregulated, and cell cycle genes downregulated, indications of no mb, C2-DPC and C6-DPC capacity to maturation in direct reprogramming **(Fig. 3H, Table EV4).**

In addition to C6-DPC inducing sarcomeric and cytoskeletal assembly genes such as LMOD3, and SNTB1, together with mitochondrial and oxidative metabolism components including COX6A2 and ACADVL **(Fig. 3H, EV3B, Table EV4)**, we identified a new category of affected genes: C6-DPC specifically suppresses the interferon/inflammatory barrier (IFIT1, IFIT3, IFI44L, CXCL11) **(Fig. 3H, Table EV4)** . Some inflammatory and NF-κB-associated genes (such as IL1A, IL1B, IL24, IL32, CSF2, CSF3, CXCL8, NFKBIA, NFKB2, STAP2, and TNFRSF21) remained significantly upregulated under C6 conditions, indicating that myogenic reprogramming occurs alongside a persistent inflammatory transcriptional program **(Fig. 3H, Table EV4)**. C6-DPC does not globally suppress stress-associated programs, instead, biased FGFR signaling combined with ALK1 and TGFBR2 inhibition selectively enables myogenic structural and metabolic maturation to dominate over fate-stabilizing fibroblast programs. These findings support a model in which efficient lineage conversion is achieved not by erasing inflammatory identity, but by functionally overriding it through targeted signaling bias that unlocks irreversible myogenic commitment. To confirm the metabolic remodeling, we show that C6-DPC induced increased mitochondrial Mito-orange uptake indicating improved bioenergetic capacity **(Figs. 3I-J)**. Further, prolonged C6-DPC treated cells exhibited pronounced sarcomere organization and myofiber maturation (**Fig. 3K)**. Together, these findings identify C6-DPC-mediated FGFR1c-biased signaling as a key determinant of functional myogenic reprogramming.

### Inflammatory signaling limits myogenic conversion

To investigate potential barriers to myogenic reprogramming, we performed single-nucleus RNA sequencing of nuclei isolated from the remaining mononucleated cell population following MYOD induction, with or without C6-DPC treatment (Fig. 4A). Within this population, C6-DPC-treated cells exhibited elevated expression of TNFRSF19, TNFRSF21, TDO2, PLAT, and matrix metalloproteinase (MMP) genes, consistent with a transcriptional program associated with TNF-related signaling, stress adaptation, and extracellular matrix remodeling (**Fig. EV3D**). Inflammatory gene-expression scores also varied across the identified cell populations, linking inflammatory programs to distinct cell states during myogenic conversion (**Fig. 4B, EV3C**).

**Figure 4.**
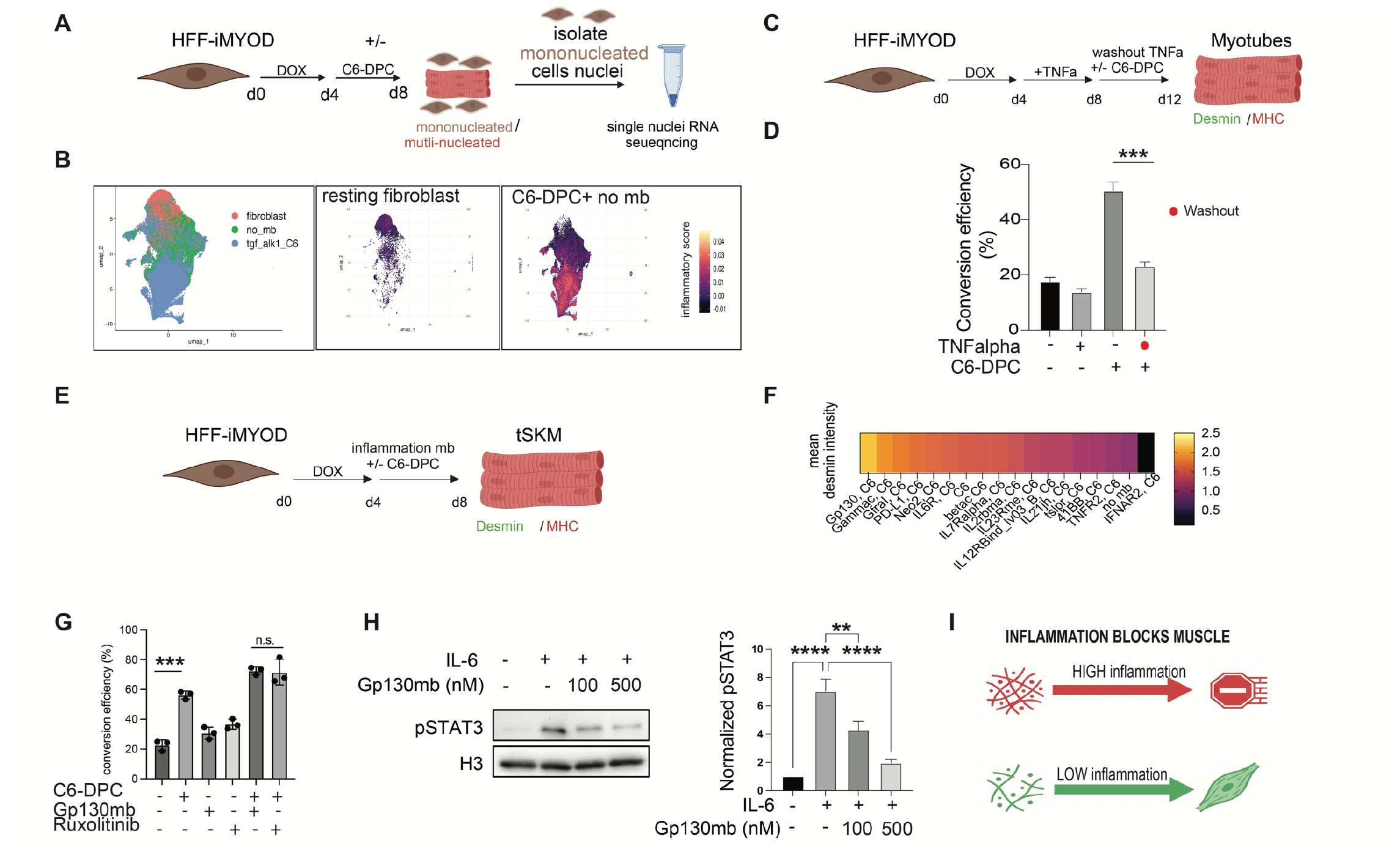
Inflammation blocks muscle reprogramming. (A) Schematic showing single nuclei RNA sequencing of original resting fibroblast, day 8 tSKM no mb and day 8 tSKM C6-DPC treated mononucleated cells. (B) UMAP showing all the single nuclei RNA sequenced. UMAP showing the fibroblast (left) and, no mb and C6-DPC (combined, right) inflammation score based on inflammatory cytokines and receptor expression level. (C) HFF-iMYOD were induced with doxycycline for 4 days, followed by TNF-alpha treatment for next 4 days to induce inflammation followed by treatment of C6-DPC for 4 days. (D) Bar graph of fibroblast to muscle conversion efficiency as in indicated treatments, quantification of immunofluorescence intensity of desmin and DAPI (n=3). (E) Schematic of fibroblast to muscles conversion, where HFF-iMYOD were induced with doxycycline for 4 days, followed by treatment of C6-DPC and inflammatory minibinders for another 4 days. (F) Heatmap for mean desmin intensity fold change of two independent minibinder screens. (G) Bar graph of fibroblast to muscle conversion efficiency in C6-DPC after treatment with C6-DPC, gp130mb, Ruxolitinib, C6-DPC+gp130mb, C6-DPC+ Ruxolitinib treatments between day4 to day8 of reprogramming(n=3). (H) Representative immunoblot of HFF cells treated with indicated compounds. Graphs showing normalized pSTAT3 signal (n=5). (I) Schematic model showing the conversion of fibroblast to skeletal muscles requires low inflammatory signal. Data are presented as mean ± SD with two-tailed unpaired Student’s *t*-tests. Where, *ns* for not significant \**p* < 0.05; \*\**p* < 0.01; \*\*\**p* < 0.001 and \*\*\*\**p*<0.0001.

To test whether inflammatory stimulation impairs conversion, we exposed HFF-iMYOD cells to TNF-α and subsequently removed the cytokine prior to treatment with or without C6-DPC (Fig. 4C; Zhao et al., 2015). Following TNF-α washout, conversion efficiency was significantly lower in C6-DPC-treated cultures previously exposed to TNF-α than in those without prior exposure (**Fig. 4D**), indicating that transient inflammatory stimulation can have lasting effects on myogenic conversion. Further, to determine whether inflammatory stimulation counteracts C6-DPC-enhanced myogenic conversion, we treated HFF-iMYOD cells with C6-DPC in the presence or absence of TNF-α from day 4 to day 8 following MYOD induction. Co-treated TNF-α significantly reduced conversion efficiency to 38%, These results indicate that TNF-α attenuates the enhancement of myogenic conversion induced by C6-DPC **(Fig. EV3E)**.

We next screened minibinders targeting inflammation-associated receptors for their effects on myogenesis (Fig. 4E,F, **Table EV5-6**). A gp130-targeting minibinder further increased conversion efficiency when combined with C6-DPC **(Fig. 4G, Table EV6)**. Consistent with inhibition of gp130-mediated signaling, this minibinder reduced IL-6-induced STAT3 phosphorylation at 100 and 500 nM in HFF as well as in EA.hy926 cells (**Fig. 4H**, **EV3F**). T As a pharmacological comparison, ruxolitinib suppressed IL-6-induced STAT3 phosphorylation to near-basal levels, also showed similar conversion efficiency in combination of C6-DPC in comparison to gp130mb(**Figs. 4G**, **EV3G**). These results demonstrate that the gp130 minibinder attenuates IL-6-STAT3 signaling in HFF and EA.hy926 cells. Together, these findings identify inflammatory signaling as a modifiable barrier to myogenic conversion and support targeting the IL-6-gp130-STAT3 pathway to improve the response to C6-DPC **(Fig. 4I).**

### Designed protein cocktail promotes iPSCs into muscle differentiation

To determine if the C6-DPC has a general myogenic capacity, beyond transdifferentiation, we analyzed the cocktail in iPSC derived skeletal muscle differentiation protocol. WTC11(dox inducible MyoD) human iPSCs were treated with doxycycline to initiate myogenic commitment, followed by differentiation in skeletal muscle medium and subsequently treated with C6-DPC **(Fig. 5A)**. Importantly, C6-DPC treatment resulted in robust formation of multinucleated myotubes expressing canonical muscle markers desmin and Myosin Heavy Chain (MHC) (**Fig. 5B**). Quantitative analysis revealed that C6-DPC treatment significantly increased myotube area coverage relative to control conditions lacking minibinders and the mean fluorescence intensity of MHC was significantly elevated in C6-DPC treated cultures **(Figs. 5C-D)**. These data indicate that C6-DPC increases sarcomeric protein accumulation and myofiber maturation during iPSC derived muscle differentiation.

**Figure 5.**
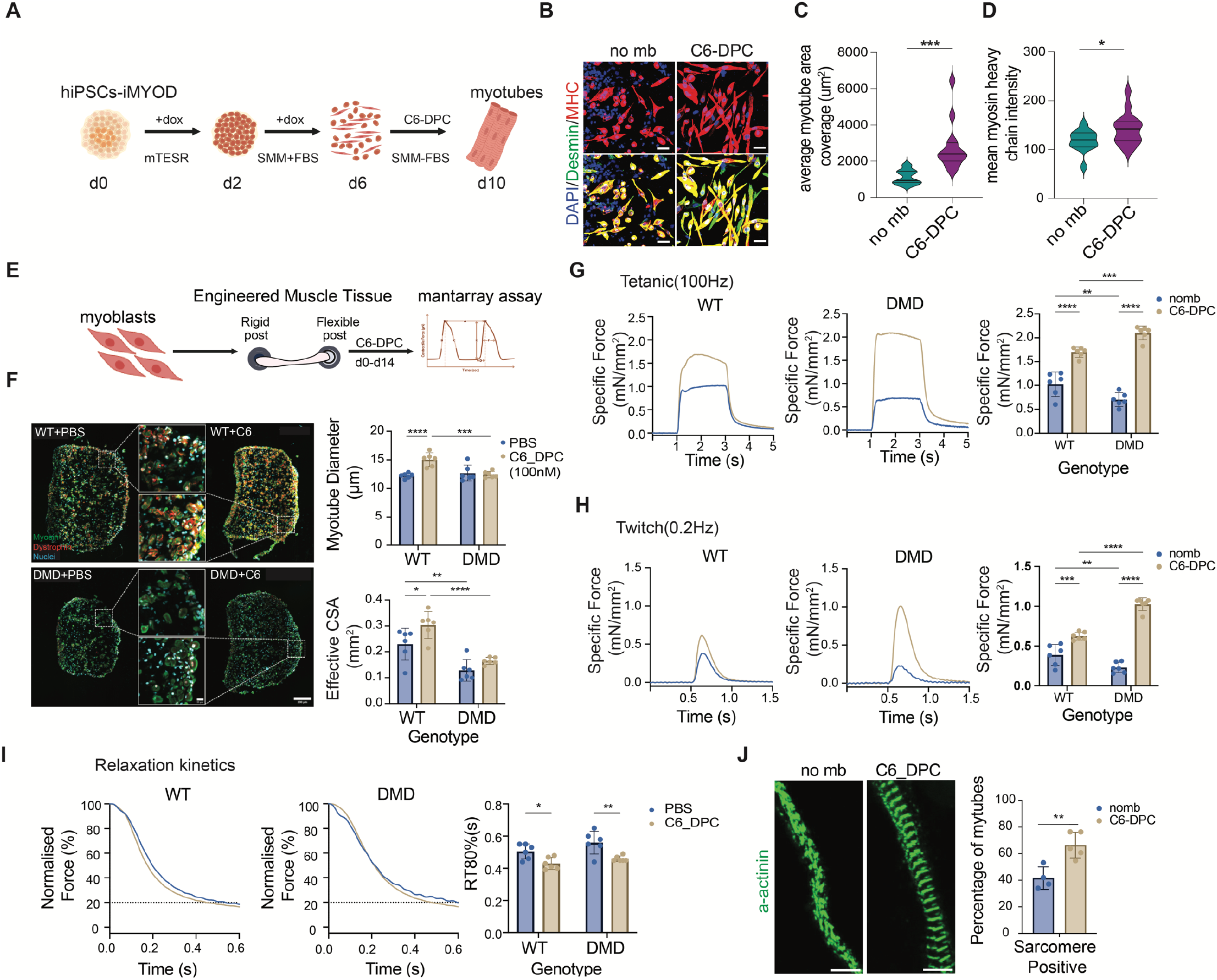
C6-DPC treatment enhances iPSCs differentiation into skeletal muscles, enhances contractile force and accelerates kinetics in iPSC-derived EMTs. (A) Schematic showing iPSCs-iMYOD differentiation into myotubes in C6-DPC for 4 days, fixed and stained for muscle makers. (B) Representative images of iPSCs differentiated into myotubes upon no mb and C6-DPC treatment are stained for DAPI (blue), desmin (green) and MHC (red). Scale bar, 10um. (C) Violin plot showing average myotube area coverage upon no mb or C6-DPC treatment (n=_3). (D) Violin plot of MHC mean intensity upon no mb or C6-DPC treatment (n=3). (E) Schematic showing mantarray assay measuring the contractile forces where the 3D muscle tissue (EMTs) is scaffolded around a rigid and a flexible post. The contractile force generated by the tissue is measured with the displacement of a magnet in the flexible post. (F) Representative immunofluorescence cross-sections of WT and DMD engineered muscle tissues (EMTs) treated with PBS (vehicle) or C6-DPC (100 nM). Sections are stained for myosin heavy chain (green), dystrophin (red), and nuclei (blue). Scale bar: 200 µm. Insets show higher magnification of myotube architecture (Scale bar: 20 µm). Graphs show quantification of average myotube diameter (Top) and cross sectional area (Bottom) (n=6 ). (G) Representative tetanic (100Hz) force traces (left) and quantification of specific tetanic force (right), normalized to effective CSA (n=6 ). (H) Representative twitch (0.2Hz) force traces (left) and quantification of specific twitch force (right), normalized to effective CSA (n=6 ). (I) Normalized relaxation traces (left) and quantification of the time to 80% relaxation (RT80%) (right), indicating improved relaxation kinetics with C6-DPC treatment (n=6 ). (J) Representative immunofluorescence of longitudinal image of WT Engineered muscle tissue showing sarcomeric organisation after treatment with no mb and C6-DPC stained with Alpha-actinin antibody. Scale bar, 5 um. Graph represent percentage of sarcomere positive myotubes (nomb n=4, C6-DPC n=5 ). Data are presented as mean ± S.E.M. *P < 0.05, **P < 0.01, ***P < 0.001, ****P < 0.0001 by two-way ANOVA with Tukey’s multiple comparisons test.

### C6-DPC enhances structural and functional maturation of iPSC-derived engineered muscle tissues

To determine whether C6-DPC induces functional gains in a 3D muscle tissue context, we generated engineered muscle tissues (EMTs) from both wild-type and dystrophin-deficient (DMD) iPSC derived myoblasts in the presence or absence of C6-DPC and assessed contractile function using the Mantarray magnetometric platform after 14 days of culture (**Fig. 5E**). Immunofluorescence cross-sections of C6-DPC-treated EMTs revealed denser, more organized myofiber architecture compared to PBS-treated controls in both genotypes (**Fig. 5F**). We quantified this structural maturation, observing a significant increase in effective cross-sectional area with C6-DPC treatment in wild-type tissues (PBS: 0.229 vs C6-DPC: 0.304 mm²), with a smaller, non-significant increase in DMD EMTs (PBS: 0.128 vs C6-DPC: 0.165 mm²) (Fig. 5F). . C6-DPC treatment also increased mean myotube diameter in WT tissues (PBS: 12.24 vs C6-DPC: 15.06 µm), though this hypertrophic effect was not significant in DMD EMTs (**Fig. 5F**), indicating a blunted hypertrophic response in dystrophin-deficient myotubes.

Critically, these structural gains translated directly into enhanced intrinsic contractile output. To accurately assess muscle quality independent of the larger tissue cross-section, we evaluated specific force by normalizing the active force to the effective CSA. Under tetanic stimulation (100 Hz), C6-DPC treated EMTs exhibited significantly elevated specific force, generating approximately 1.6-fold greater than control force in WT tissues (PBS: 1.03 vs C6-DPC: 1.69 mN/mm²) and a profound 3 fold increase in DMD tissues (PBS: 0.70 vs C6: 2.10 mN/mm²) (**Fig. 5G**). Single twitch responses (0.2 Hz) followed a similar pattern, C6-DPC treatment increased specific force by ∼1.6 fold in WT and ∼4.5 fold in DMD EMTs specific twitch force (**Fig. 5H**). This demonstrates that the contractile improvement exceeds what can be attributed to increased tissue size alone, indicating that C6-DPC treatment fundamentally enhances the force-generating capacity of the myofibers themselves.

To further assess functional maturation, we examined relaxation kinetics following tetanic contraction. C6-DPC treatment significantly accelerated the time to 80% force relaxation (RT80%) in both WT (PBS: 0.505s vs C6-DPC: 0.430s, Δ75 ms) and DMD tissues (PBS: 0.560s vs C6-DPC: 0.460s, Δ100 ms) (**Fig. 5I**). Faster relaxation is a hallmark of mature skeletal muscle, reflecting efficient calcium reuptake and cross-bridge detachment kinetics. Additionally, α-actinin staining revealed an increased proportion of myotubes with organized sarcomeres following C6-DPC treatment (**Fig. 5J)**. Together, these results demonstrate that C6-DPC enhances structural maturation and contractile performance in the skeletal-muscle models examined.

Collectively, these results demonstrate that C6-DPC does not simply produce larger muscle; it produces muscle with intrinsically greater force. The convergence of increased specific force, accelerated relaxation kinetics, and enhanced myofiber fusion defines a coordinated maturation program that is preserved across both healthy and dystrophin-deficient backgrounds, establishing synthetic receptor-specific signaling as a viable strategy for functional muscle regeneration.

### Cooperative regulation of promoter accessibility enhanced during C6 cocktail induced myogenic reprogramming

Bulk RNA sequencing revealed that treatment with the designed protein cocktail (C6-DPC) induces a robust transcriptional program consistent with myogenic differentiation and muscle maturation in transdifferentiated skeletal muscle (tSKM) cells **(Figs. 6A-B)**. The C6-DPC treatment significantly upregulated genes associated with muscle progenitor activation, cytoskeletal remodeling, and paracrine signaling, including MSX1, HGF, SVIL, FIBCD1, SNAI1, CMKLR1, PTGS2, and PGF, reflecting a coordinated shift toward a pro-myogenic and growth-permissive state **(Figs. 6B, Table EV7).** KEGG analysis, Gene Ontology enrichment and reactome analysis revealed activation of PI3K-AKT, MAPK/ERK, JAK-STAT, calcium, VEGF, Wnt signaling, and arachidonic acid metabolism **(Figs. EV4A-C)**, collectively indicating enhanced growth factor responsiveness, survival signaling, and metabolic remodeling downstream of C6-DPC treatment. Notably, induction of PTGS2 and PTEGS by C6-DPC **(Fig. EV4D)**, aligns with established roles of prostaglandin signaling in muscle growth and regeneration (Ho et al., 2017; Wang et al., 2025). The upregulation of HGF and the myokine FIBCD1 (**Fig. 6B)** further suggests activation of autocrine and paracrine programs that support myofiber growth and tissue remodeling (Tatsumi et al., 1998; Graça et al., 2022), consistent with the increased myotube dimensions observed following C6-DPC treatment. Developmental regulators such as NR4A2, ID3 and SNAI1 **(Fig. 6B, Table EV7)**, that are associated with early lineage plasticity and morphogenetic programs, were also elevated (Benezra et al., 1990; Soleimani et al., 2012), suggesting partial reactivation of developmental transcriptional states that may synergize with chromatin remodeling to enhance maturation **(Fig. EV4E)**.

**Figure 6.**
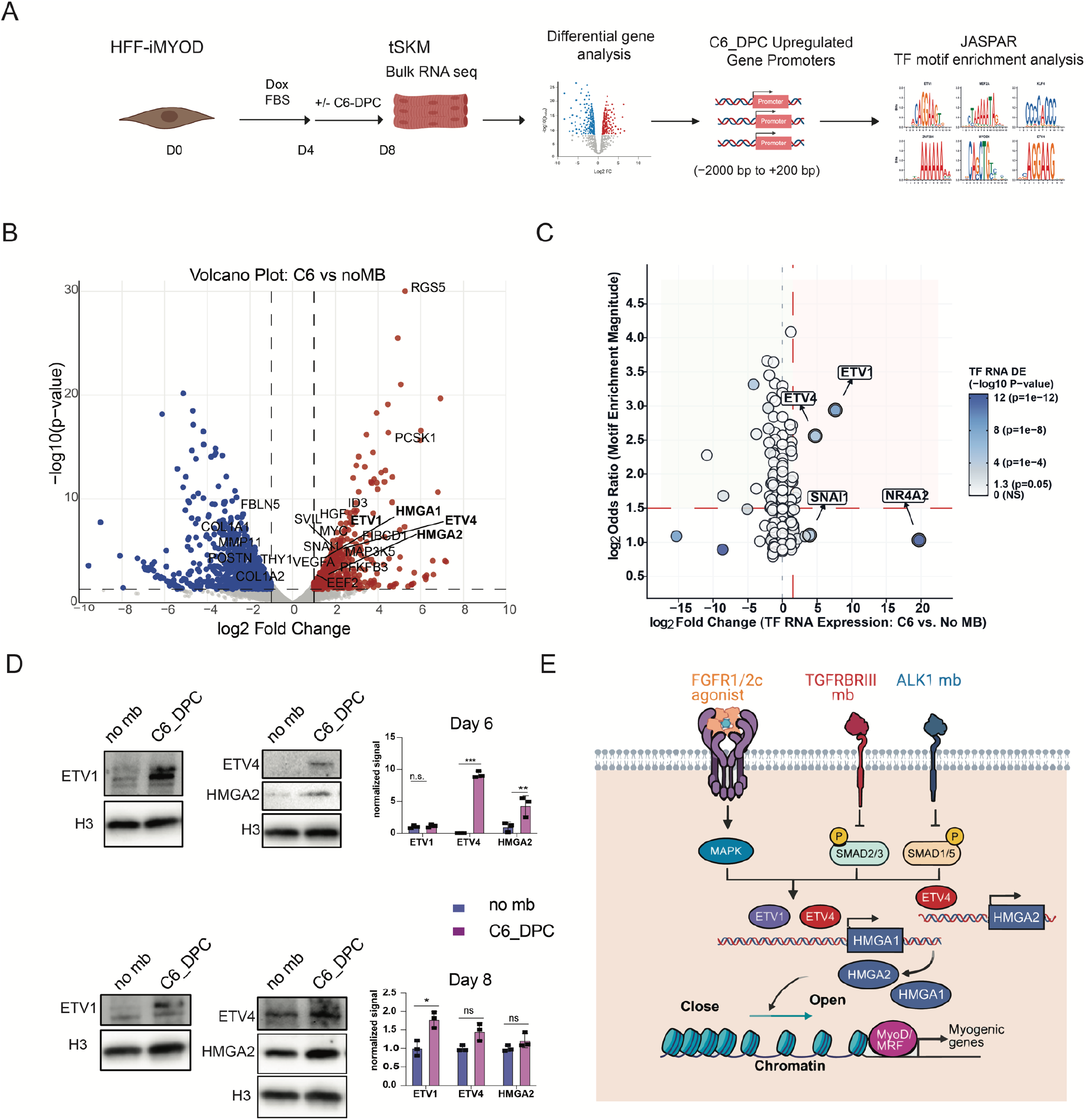
C6-DPC mediates promyogenic chromatin remodeling. (A) Experimental workflow for bulk RNA sequencing and transcription factor motif enrichment analysis. HFF-iMYOD cells treatment with or without C6-DPC from day 4 to day 8. Promoters of genes upregulated by C6-DPC identified by differential gene analysis, where the upregulated genes promoters are extracted as −2,000 to +200 bp relative to the transcription start site, were analyzed for enrichment of transcription factor motifs using JASPAR. (B) Volcano plot showing differential gene expression between C6-DPC-treated and no-minibinder (noMB) control cells at day 8. Red and blue points indicate significantly upregulated and downregulated genes, respectively (P < 0.05 and |log₂ fold change| ≥ 1); grey points indicate genes below these thresholds. Selected genes are labeled. (C) Integration of promoter motif enrichment with expression changes of the corresponding transcription factors. The x-axis indicates transcription factor RNA log₂ fold change in C6-DPC versus noMB cells, and the y-axis indicates log₂ odds ratios for motif enrichment in promoters of C6-DPC-upregulated genes. Point color represents the −log₁₀ P value for differential transcription factor expression. ETV1, ETV4, SNAI1, and NR4A2 are highlighted which are C6-DPC -induced drivers. Significance calculated by one-sided Fisher’s exact test with Benjamini– Hochberg procedure correction (RNA-seq data: GEO GSE352439). (D) Western blot showing ETV1, ETV4 and HMGA2 protein levels at day6 and day 8 of reprogramming where no mb and C6-DPC treatments are compared. Quantification of immunoblot analysis of the ETV1/ETV4/HMGA2 on day 6 and day 8 of reprogramming (n=3). Data are presented as mean ± SD with two-tailed unpaired Student’s *t*-tests. Where, *ns* for not significant \**p* < 0.05; \*\**p* < 0.01; \*\*\**p* < 0.001 and \*\*\*\**p*<0.0001. (E) We propose a model where C6-DPC mediates enhanced myogenic reprogramming through cooperative transactivation activity of ETV1 and ETV4 on HMGA1 promoter and ETV4 activity on HMGA2 promoter. HMGA1 and HMGA2 mediated chromatin opening, enables further activation of myogenic gene expression.

To predict transcriptional regulators underlying enhanced myogenic reprogramming induced by the FGFR1/2-c-agonist based C6 designed protein cocktail (C6-DPC), we performed bulk RNA sequencing followed by motif enrichment analysis of the promoters of differentially upregulated genes relative to the no-minibinder control. Promoters were defined as regions spanning −2,000 to +200 bp relative to the transcription start site **(Fig. 6A,B)**. Enrichment analysis identified prominent zinc-finger/KLF and ETS-family motifs, together with myogenic MEF2 motifs **(Fig. 6C).** ETV1 and ETV4 motifs were enriched in 45.4% and 52.4% of foreground promoters, respectively (**Fig. 6C, Table EV8)**. Notably, ETV1 and ETV4 were also among the differentially upregulated genes, identifying them as candidate mediators of the C6-DPC. Interestingly, ETV1 and ETV4 have predicted binding sites at HMGA1 distal promoters and ETV4 has a binding site on HMGA2 (**Table EV9,10)**.

Protein expression measurements revealed ETV4 and HMGA2 increased significantly at day 6 of reprogramming, whereas ETV1 showed no significant change **(Fig. 6D)**. At day 8, ETV1 increased significantly, while ETV4 and HMGA2 did not differ significantly from controls **(Fig. 6D).** These findings support a proposed model in which FGFR1/2-c stimulation, combined with TGFBRII and ALK1 antagonism, engages ETS-associated transcriptional programs and HMGA-dependent chromatin regulation to facilitate MYOD-driven myogenic gene expression (**Fig. 6E**). Direct ETS binding and its contribution to chromatin remodeling remain to be established.

We propose a model in which AI-designed receptor-modulatory minibinders and inflammation inhibitor minibinders rewire extracellular signaling to overcome barriers to myogenic reprogramming. In this framework, an engineered FGFR1/2-c-specific agonist activates pro-myogenic MAPK/ERK signaling, leading to induction of ETS transcription factors that promote muscle gene expression and metabolic maturation. In parallel, inhibitory minibinders targeting TGFβ signaling components, including ALK1 and TGFBR2, suppress SMAD2/3-driven fibrotic and anti-myogenic programs that otherwise divert cells toward non-muscle lineages. Furthermore, inflammation inhibition by designed minibinders is beneficial for muscle maturation since TNF-α signaling disrupts activation of the MyoD-myogenic feed-forward network, and gp130-IL–STAT3 signaling reinforces a competing fibro-inflammatory transcriptional state that overrides differentiation enforcing a non-myogenic cell state (Guttridge et al., 2000; Serrano et al., 2008; Tierney et al., 2014).

The combined action of these signals creates a permissive transcriptional and chromatin landscape, enabling efficient MyoD driven conversion of fibroblasts into multinucleated myotubes. This rewiring of the signaling network does not only enhance structural differentiation, evidenced by increased fusion and sarcomeric organization but also promotes functional maturation, including elevated mitochondrial activity and improved contractile force in engineered muscle tissues **(Fig. 7)**. Notably, in dystrophin-deficient contexts, this approach enhances muscle performance despite the underlying genetic defect, highlighting the therapeutic potential of designed extracellular signaling to drive regenerative outcomes **(Fig. 7)**.

**Figure 7.**
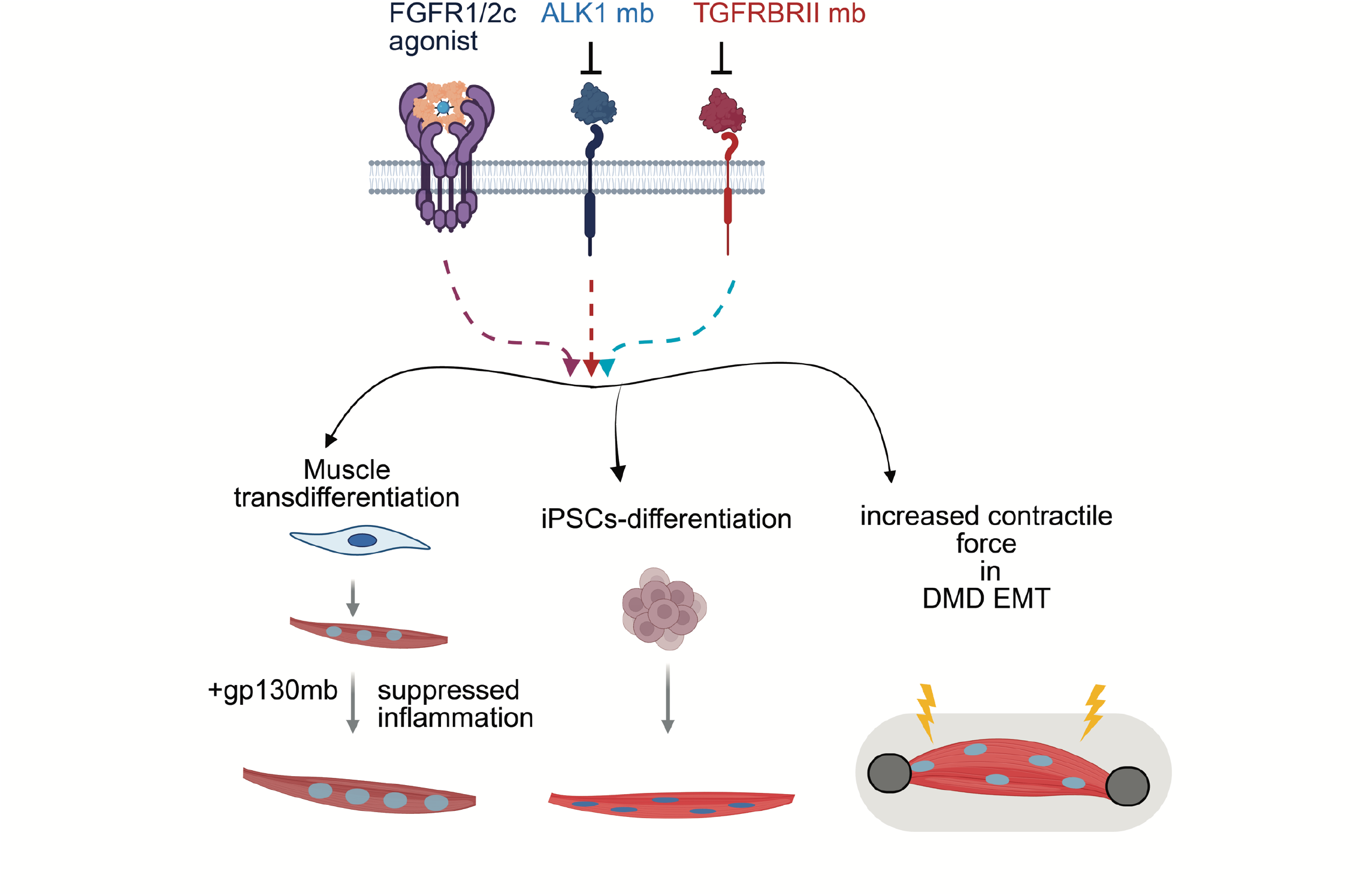
Promyogenic designed protein cocktail. A schematic showing enhanced muscle transdifferentiation, iPSC derived muscle differentiation and increased contractile forces in the 3D EMTs formed from the DMD-iPSC derived myoblast in presence of the C6-DPC.

## Discussion

Our study demonstrates that programmable extracellular signaling using AI-designed synthetic minibinders can overcome key barriers that restrict myogenic transdifferentiation. By defining an optimized reprogramming window and applying single-cell transcriptomic analysis, we identified that the majority of fibroblasts resist conversion through a coordinated barrier network involving extracellular matrix reinforcement, fibrotic lineage commitment, and transcriptional inhibitors of myogenesis. This barrier landscape reveals that inefficient reprogramming is not a stochastic process but rather reflects a structured regulatory program that stabilizes fibroblast identity. Targeted manipulation of this barrier network using AI-designed receptor modulators proved sufficient to unlock productive lineage conversion. The superior performance of the C6-DPC combining FGFR1/2-c agonism with inhibition of ALK1 and TGFBR2 signaling, demonstrates that efficient myogenic reprogramming requires simultaneous activation of pro-myogenic pathways and suppression of fate-stabilizing signaling. Unlike natural growth factors, which often trigger pleiotropic or proliferative responses, synthetic C6-DPC enables receptor isoform specific control of signaling architecture. This precision allows selective attenuation of fibrotic signaling programs while reinforcing metabolic and structural pathways associated with muscle differentiation.

This work resolves a longstanding question in the field by defining the critical, targetable nodes of small-molecule mediated direct reprogramming in muscles. Prior studies identified SB431542 and LDN-193189 as effective modulators of TGFβ-family signaling; however, their lack of specificity and the breadth of the TGFβ superfamily have limited mechanistic clarity (Jeong et al.,2021). Here, we demonstrate that two AI-designed, pathway-specific inhibitors, ALK1mb and TGFβRmb, are sufficient to precisely suppress the key TGFβ-family components that constrain myogenic reprogramming. These findings move the field from correlative small-molecule effects to a defined, target-centric framework, establishing that selective inhibition of discrete signaling nodes can recapitulate-and refine-the pro-reprogramming activity of broadly acting compounds.

Our findings identify inflammatory signaling as a novel, dominant regulatory checkpoint in myogenic reprogramming. The potent inhibitory effect of TNF-α on C6-DPC–mediated conversion demonstrates that inflammatory pathways can override pro-myogenic cues, providing a mechanistic basis for impaired regenerative capacity observed in aging and muscular dystrophy. While RTK/RSTK modulation drives unprecedented levels of transdifferentiation, conversion efficiency ultimately plateaus, revealing an intrinsic barrier. Strikingly, targeted suppression of inflammatory signaling lifts this ceiling: co-treatment with a designed gp130 antagonist significantly enhances myogenic conversion beyond this limit. Mechanistically, gp130 inhibition attenuates IL–JAK–STAT3 signaling (Selander et al., 2004; Sato et al.,2024), thereby dismantling pro-inflammatory and pro-fibrotic transcriptional programs that stabilize fibroblast identity and antagonize MyoD-driven lineage conversion. This finding suggests that inflammatory stress acts as a dominant inhibitory checkpoint that can override the pro-myogenic signals provided by FGFR1/2-c activation and ALK1/TGFBRII inhibition. These findings are supported contextually with previous studies (Berkes and Tapscott, 2005; Cao et al., 2010; Guttridge et al., 2000; Langen et al., 2004; Cai et al., 2004) showing that TNF-α signaling interferes with MyoD transcriptional activity and disrupts activation of the myogenic feed-forward network, while gp130-IL6–STAT3 signaling reinforces a competing fibro-inflammatory transcriptional state that overrides differentiation enforcing a non-myogenic cell state (Guttridge et al., 2000; Serrano et al., 2008; Tierney et al., 2014). Together, these findings position inflammatory signaling as targetable constraint on reprogramming and establish its modulation as a powerful complementary strategy to improve regenerative outcomes, particularly in diseased or aged tissues.

Importantly, the effects of C6-DPC extend beyond structural differentiation to intrinsic functional maturation. In engineered muscle tissues, C6-DPC treatment increased not only absolute contractile force but also specific force. Engineered tissues frequently produce increased force output that reflects greater cell mass rather than improved myofiber quality (Rao et al., 2018). The specific force analysis presented here directly addresses this distinction, demonstrating that C6-DPC enhances intrinsic force-generating capacity per unit cross-sectional area, providing evidence that receptor-specific signaling modulation drives contractile improvement at the myofiber level, not simply tissue expansion. This is accompanied by accelerated relaxation kinetics, a functional signature of mature calcium handling and cross-bridge cycling (Racca et al., 2013). C6-DPC-treated cells exhibited enhanced myotube maturation and significantly improved twitch and tetanic force generation, including those derived from dystrophin-deficient iPSCs. This functional rescue suggests that precise control of receptor signaling can partially compensate for the contractile deficits associated with several muscle-related disorders perhaps with matured sarcomeric arrangement. While the primary focus of this work is the signaling framework rather than DMD therapeutics, the functional rescue raises the possibility that maturation-targeted interventions could complement dystrophin-restoration strategies where functional recovery often remains incomplete despite partial genetic correction (Hoffman et al., 2011; Oskoui et al., 2025).

Disease conditions such as Dystrophic muscle have elevated inflammatory and fibrotic signals that may alter receptor expression and the response to C6-DPC. We also recognize that repeated exposure to designed minibinders could elicit an immune response. Future studies should assess immunogenicity alongside tissue distribution, duration of exposure, and effects on non-muscle cells. Local delivery or transient dosing may help limit systemic exposure, but these approaches remain to be evaluated.

Together, these findings establish programmable AI-designed receptor binders as a versatile platform for rewiring receptor signaling networks to control cell fate transitions. By combining RTK and IL-pathway activation, inhibition, and targeted receptor degradation within a single design framework, AI-designed minibinders enable rational engineering of cellular signaling environments. This strategy provides a generalizable approach for overcoming lineage barriers and enhancing functional tissue regeneration, with potential applications in muscle repair, degenerative disease modeling, and precision regenerative medicine and mitigation of GLP-1 receptor agonist–associated lean mass loss. In this context, designed protein combinations could be tailored to preserve or regenerate skeletal muscle and reduce the risk of drug-associated sarcopenia.

## Materials and Methods-

### Reagents & Tools Table

### Methods & Protocols

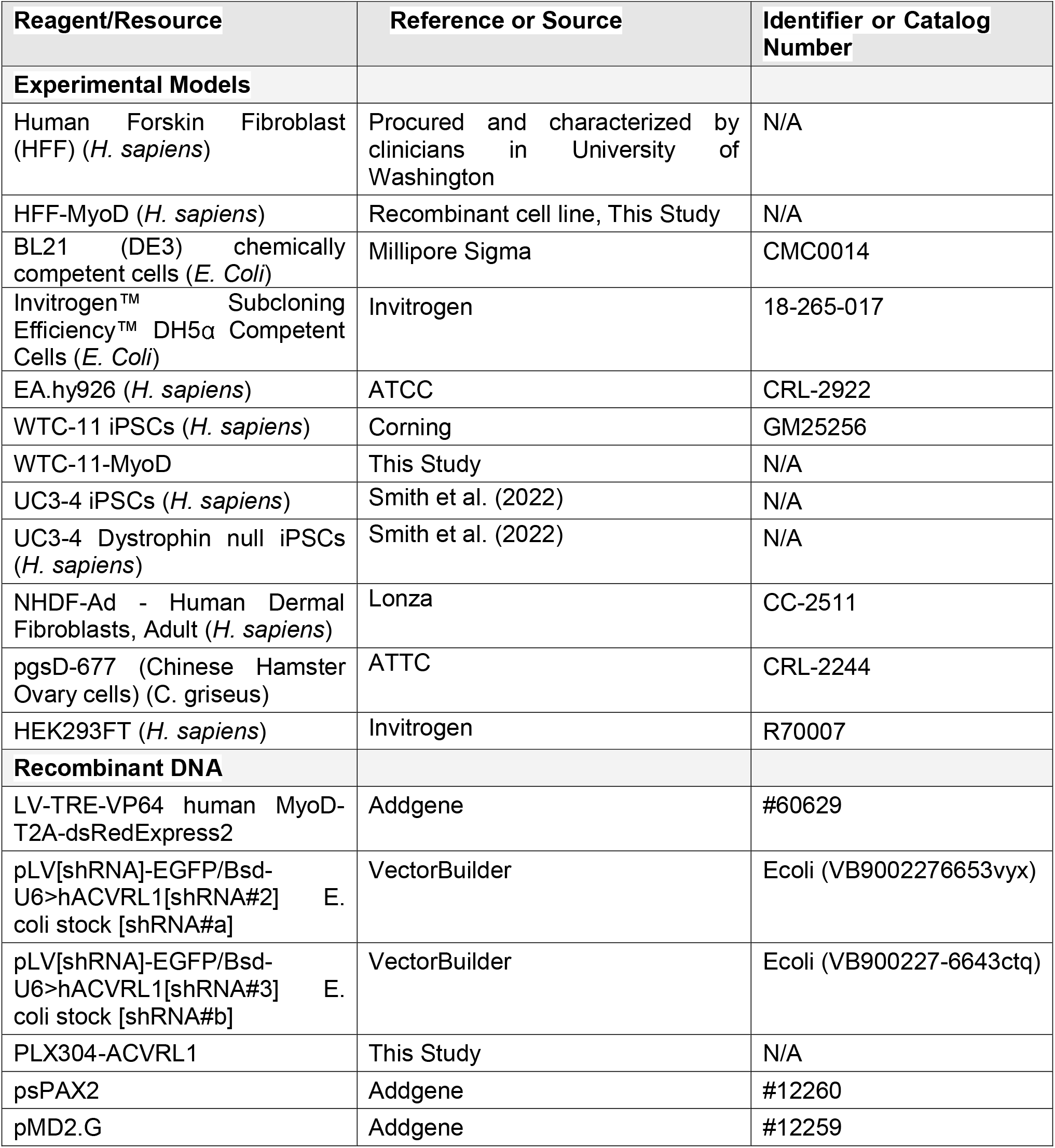
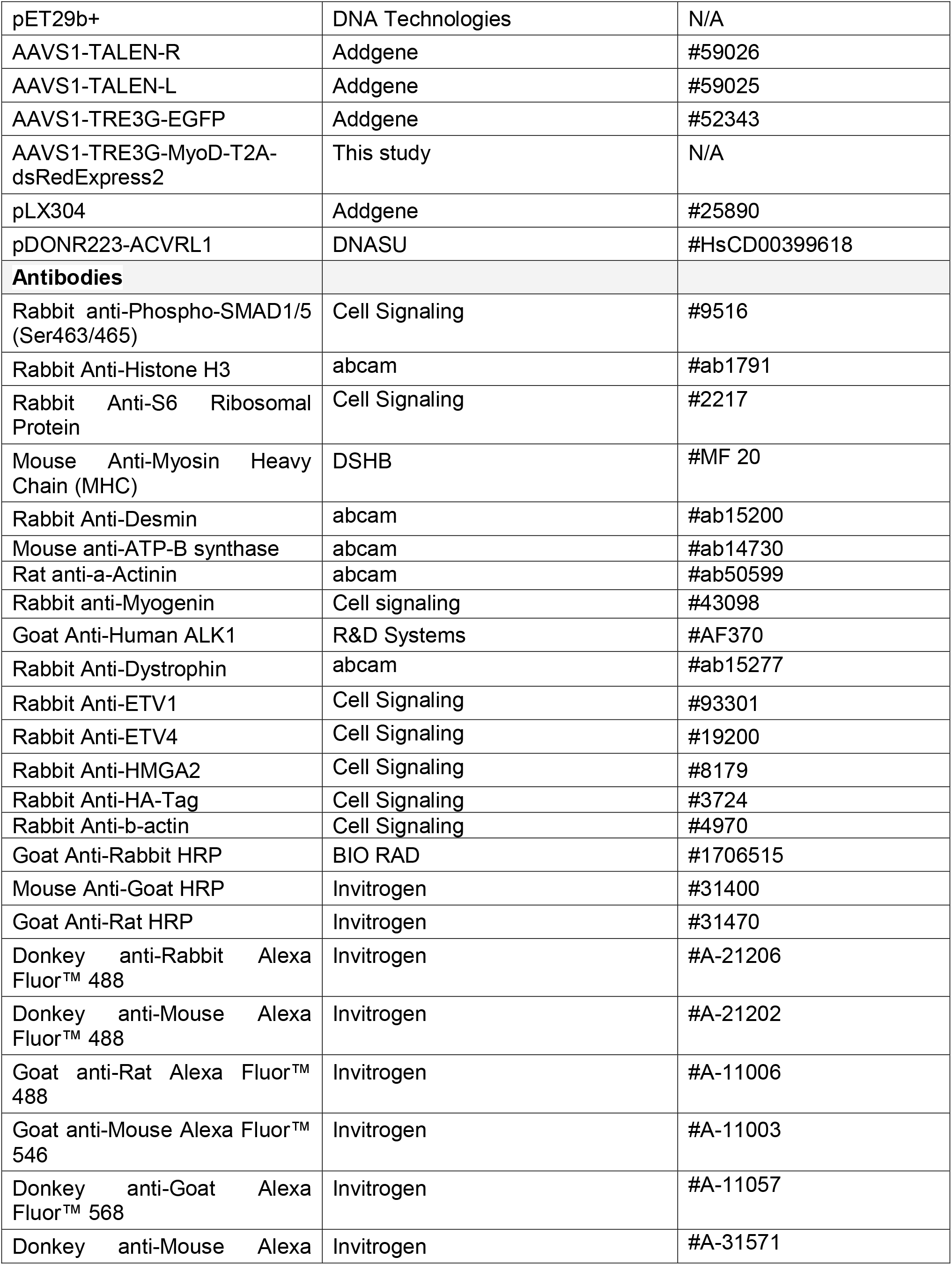
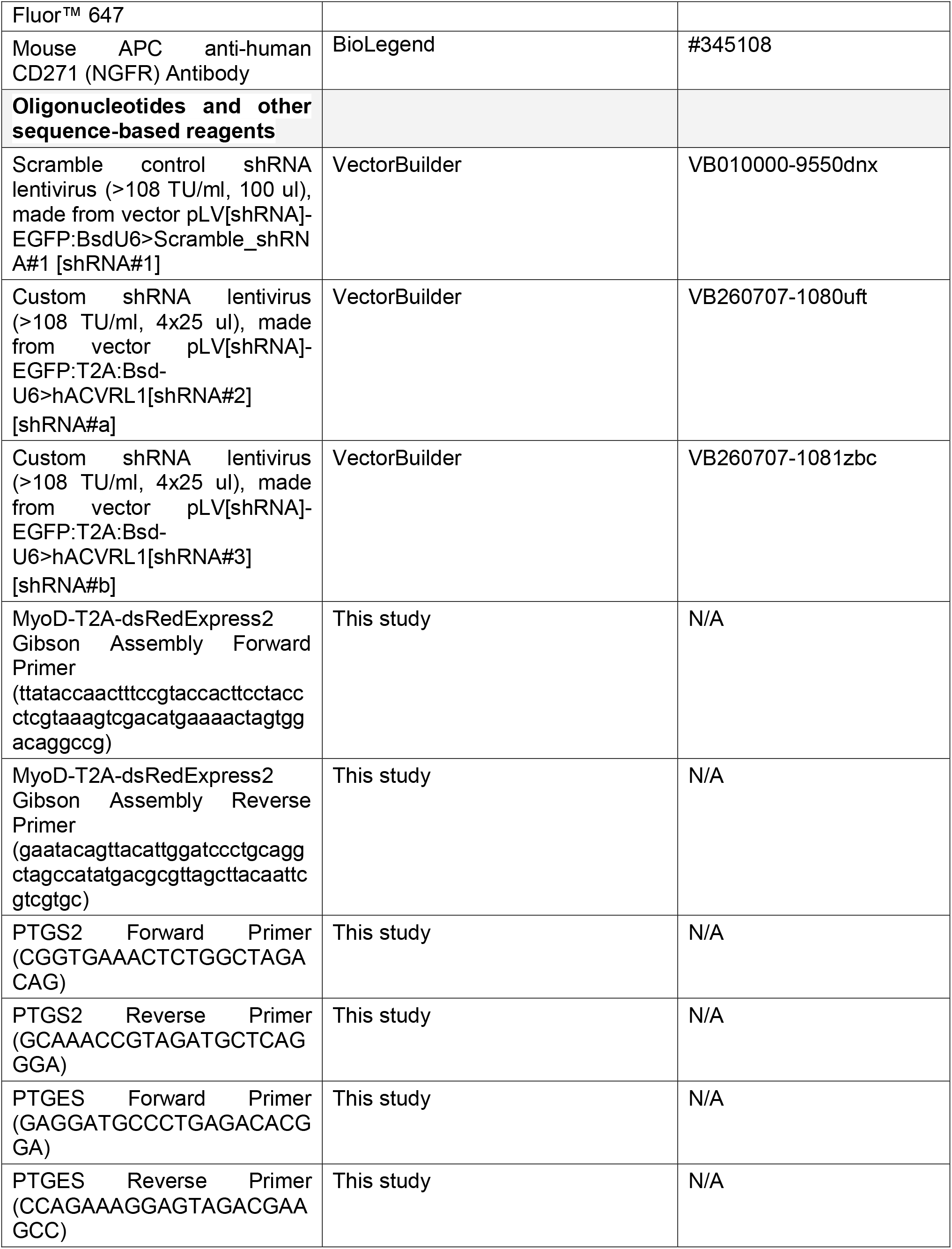
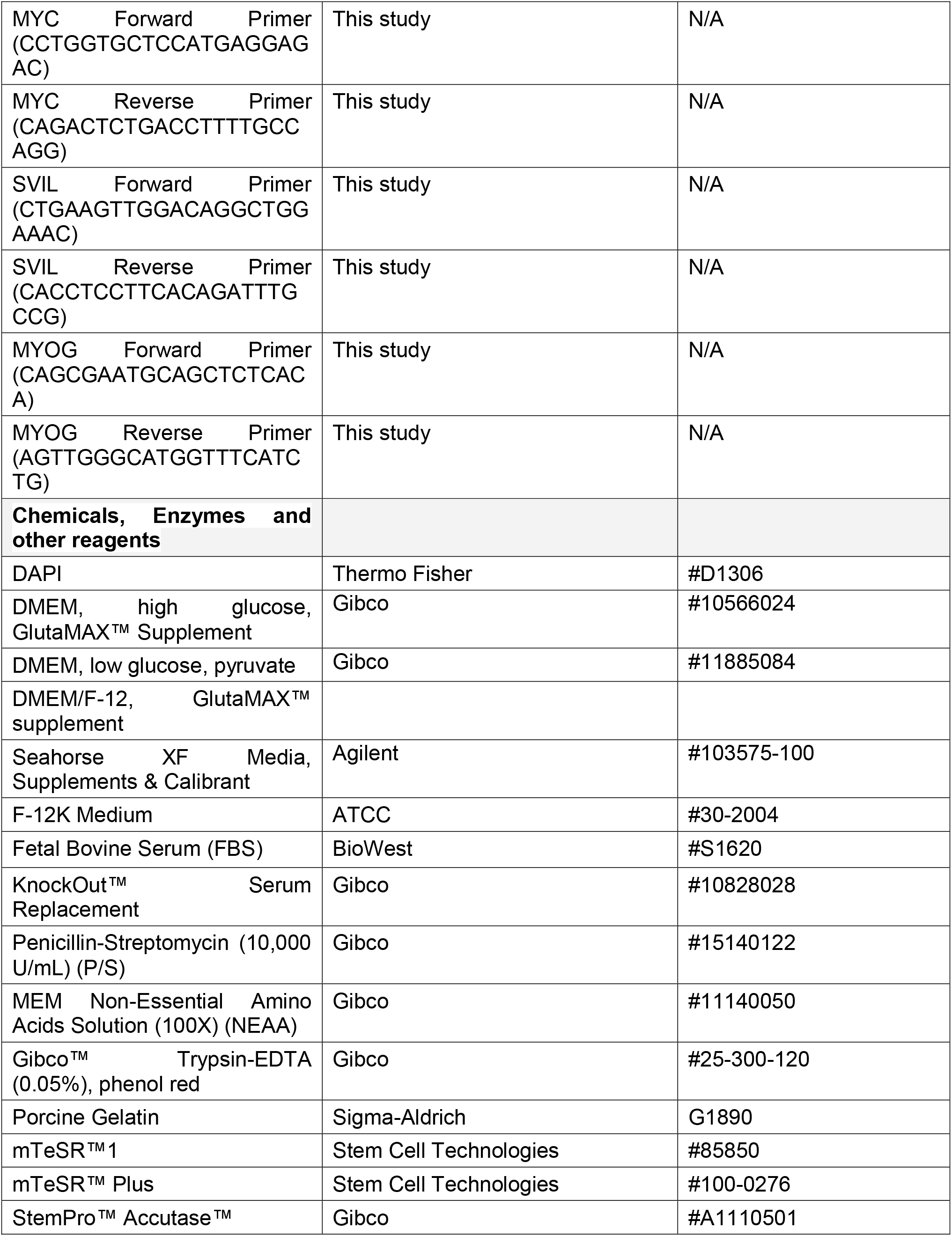
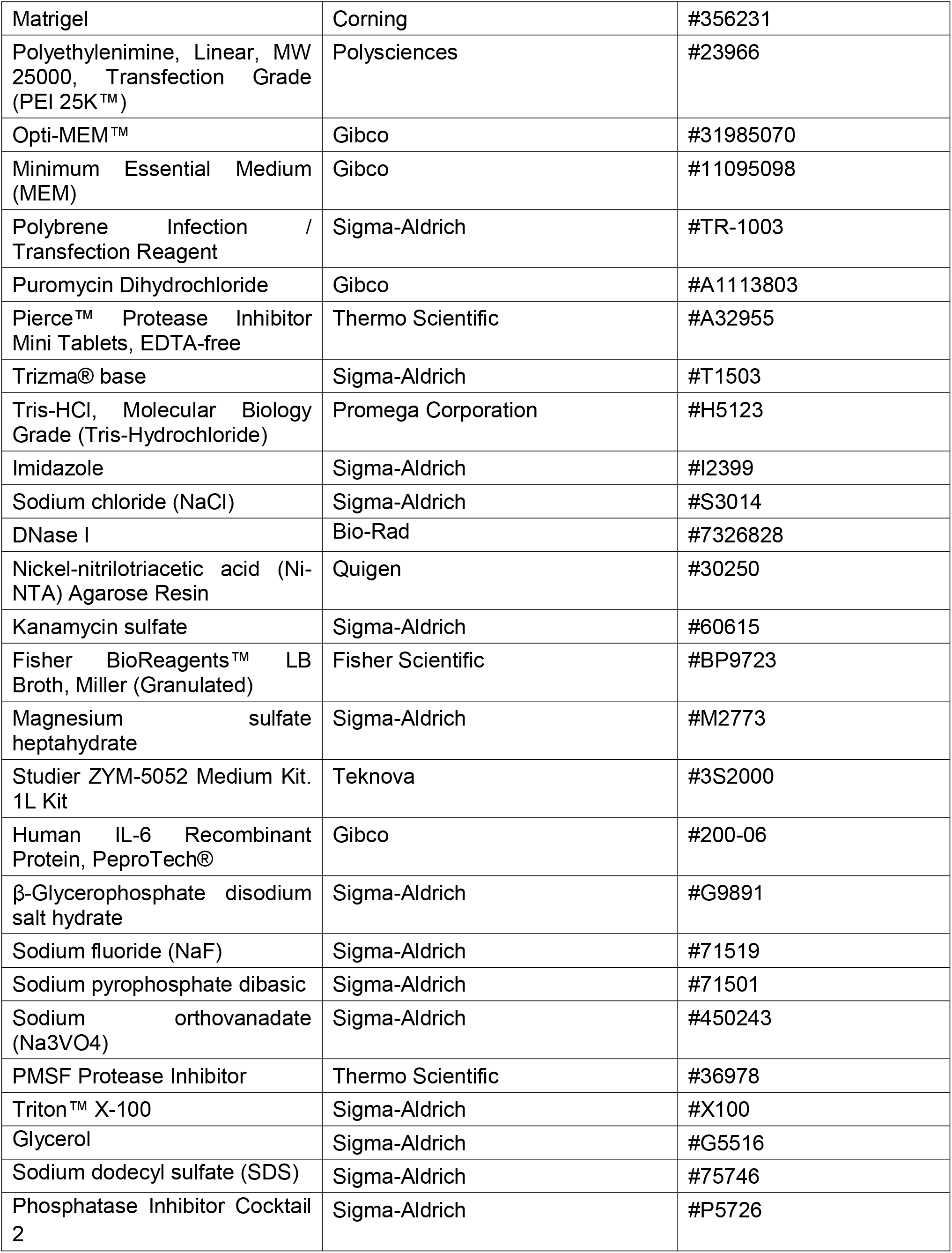
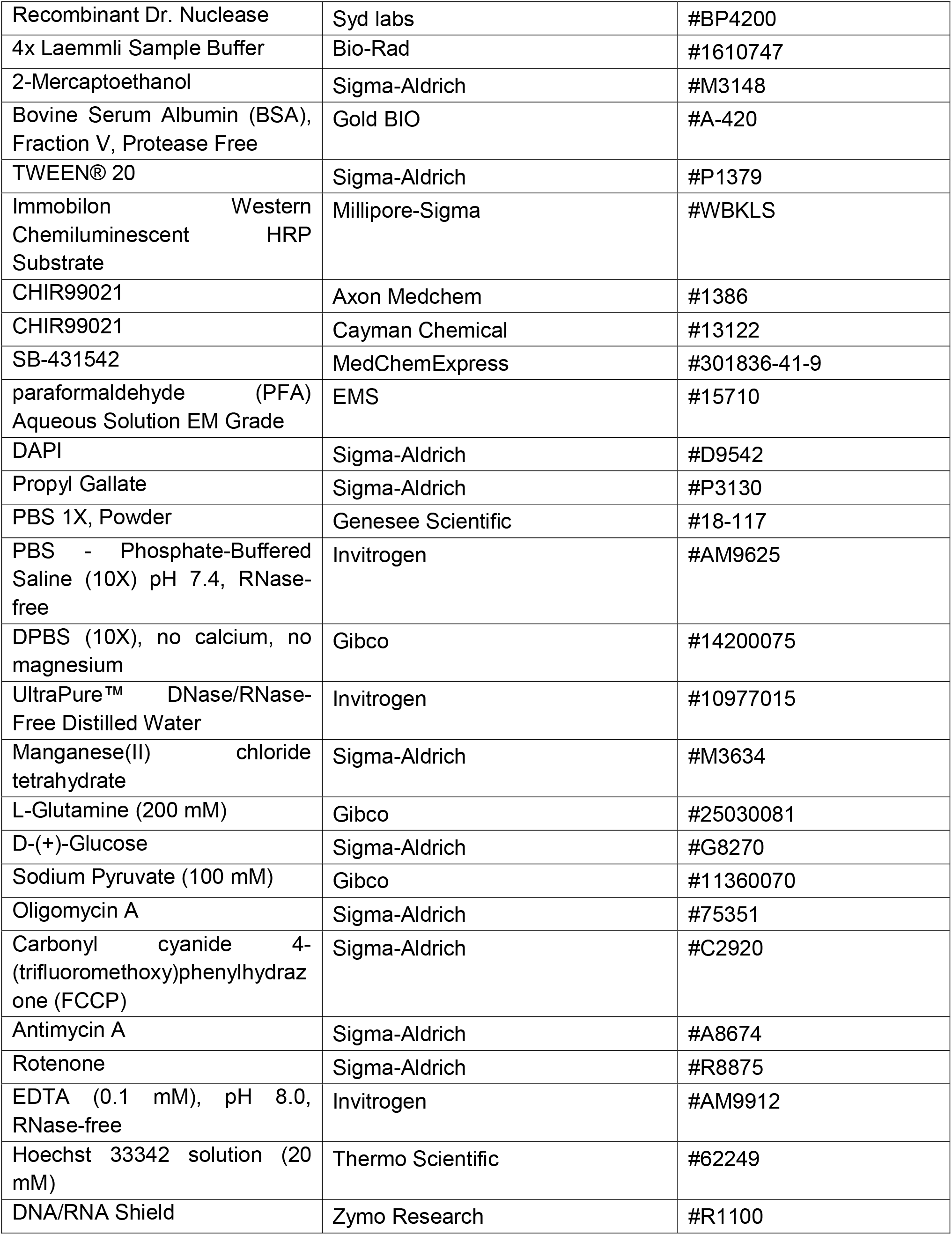
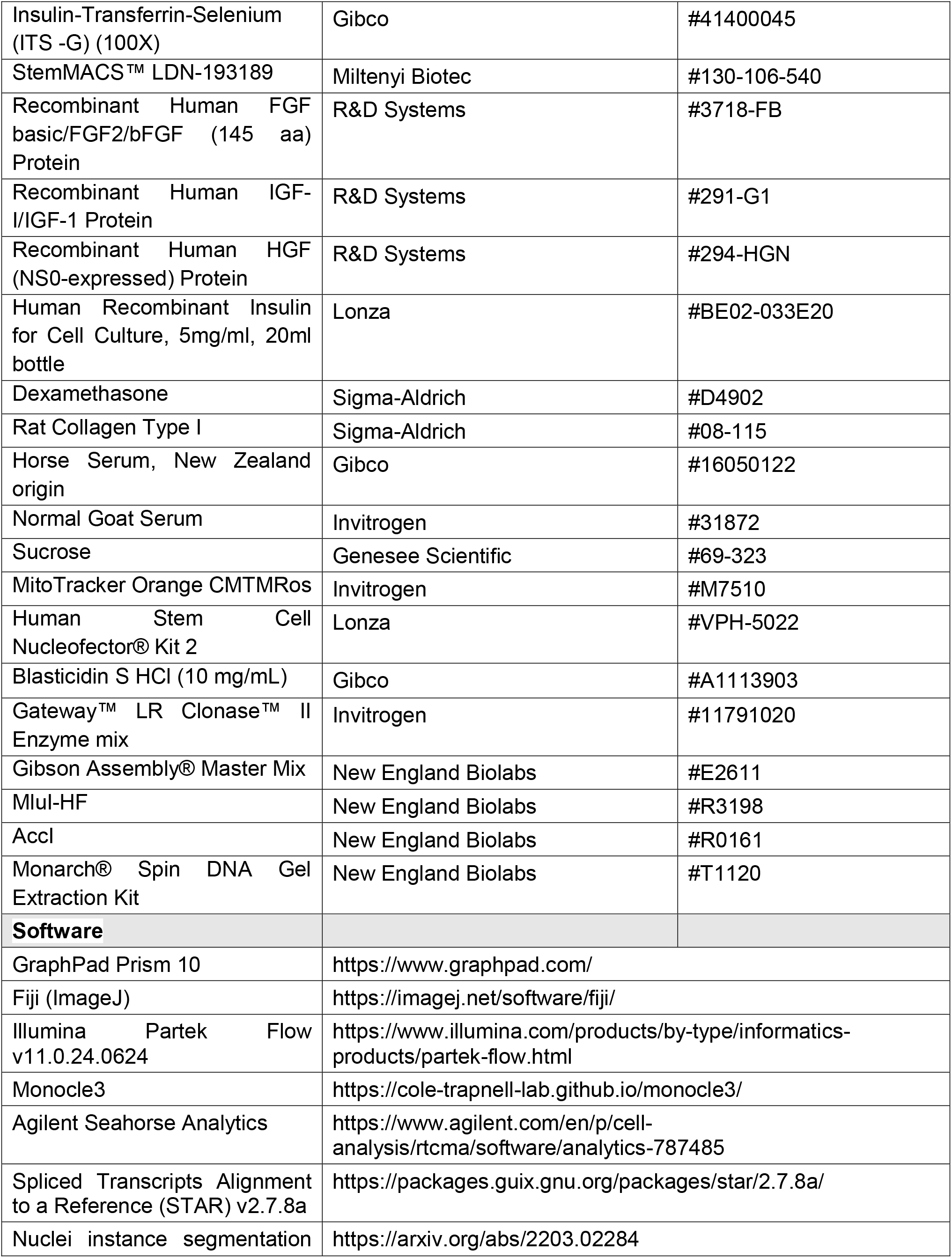
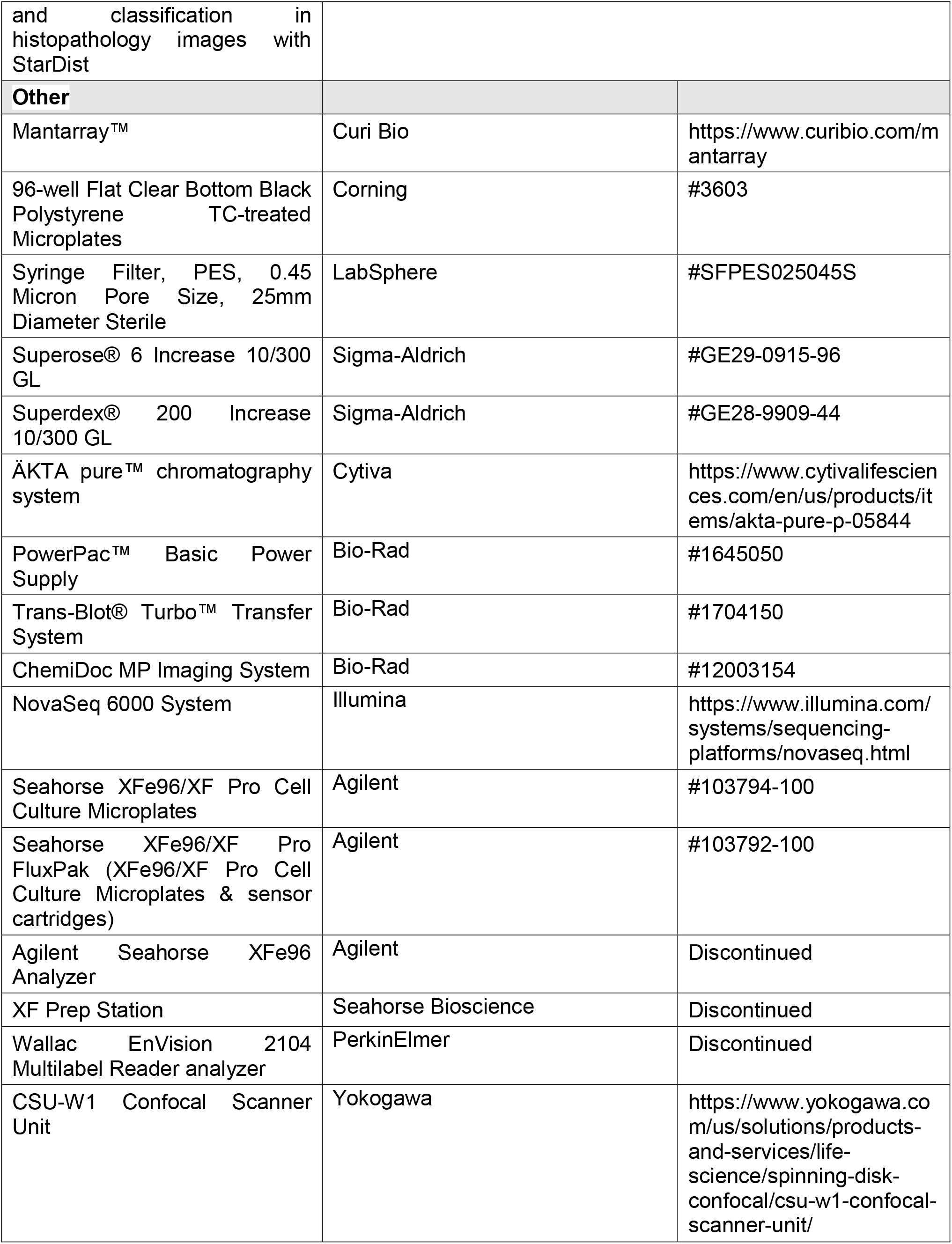
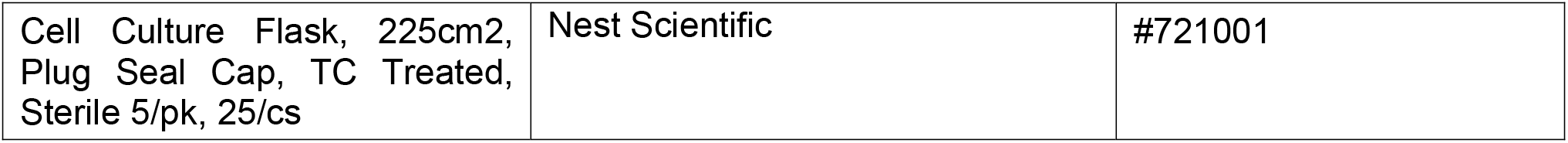

### Cell culture-CHOs, immortalized HUVECs (EA.hy926), HFF-MyoD, WTC-11-MyoD, HEK293FT

Human Foreskin Fibroblast (HFF) cells or HFF-MyoD cells were maintained in HFF Media (DMEM with high glucose & GlutaMAX™ Supplement, 10% fetal bovine serum (FBS), 1% penicillin/streptomycin (P/S), 1% NEAA). At 80% confluency, HFF-MyoD cells were passaged with Trypsin-EDTA (0.05%) and seeded at a 1:5 pellet ratio per plate with 0.1% gelatin plate coating. Chinese Hamster Ovary (CHO) cells were maintained in CHO Media (F-12K medium, 10% FBS, 1% P/S, 1% NEAA). At 90% confluency, CHO cells were passaged with Trypsin-EDTA (0.05%) and seeded at a 1:6 pellet ratio per plate. Immortalized human umbilical vein endothelial cells (immortal HUVECs, aka EA.hy926) were maintained in HFF Media. At 80% confluency, EA.hy926 cells were passaged with 0.05% Trypsin-EDTA and seeded at a 1:5 pellet ratio per plate. HEK293FT Cells were maintained in HFF Media. At 80% confluency, HEK293FT cells were passaged with 0.05% Trypsin-EDTA and seeded at a 1:12 pellet ratio per plate. Media was changed for HFF-MyoD, CHO, HEK293FT & EA.hy926 cells every other day. WTC-11-MyoD cells were maintained in mTeSR1 supplemented with 1% P/S that was changed daily. At ∼60% confluency WTC-11-MyoD were passaged with StemPro™ Accutase™ and seeded at 100K cells per 10 cm plate coated in 1:40 Matrigel solution. All cells were maintained at 37°C and 5% CO2. All the cells were authenticated by STR profiling and confirmed to be mycoplasma-negative.

### Designed protein expression and protein purification

Sequences of the designed proteins were reverse translated with optimization for Escherichia coli (E. Coli) expression, with a C-terminal glycine-serine linker followed by a 6x histidine tag. Sequences were ordered as synthetic genes from Integrated DNA Technologies within the pET29b+ vector between NdeI and XhoI cloning sites. This vector contains a kanamycin resistance marker and a T7 promoter. Plasmids were transformed into E. coli BL21 (DL3) competent cells and plated on LB with kanamycin at 50 mg/L. Transformants were inoculated into 50mL of ZYM-5052 autoinduction expression media in a 250 mL flask. Expression cultures were grown for 20 hours at 37°C with 200 rpm shaking. Cells were pelleted by centrifugation at 4000xg and resuspended in a lysis buffer consisting of 25mM Tris pH 8, 300mM NaCl, and 20mM imidazole with added PMSF protease inhibitor and DNase. Cells were lysed by sonication at 85% amplitude with 8 x 15 second pulses. Lysate was separated into soluble and insoluble fractions by centrifugation at 18,000xg. Immobilized metal affinity chromatography (IMAC) was used to purify designed protein. Nickel-nitrilotriacetic acid (Ni-NTA) resin was initially equilibrated with a 5 column volume (CV) lysis buffer. Supernatant was poured over the columns, followed by 20 CV wash buffer (25mM Tris pH 8, 400mM NaCl, 30mM imidazole). Protein was eluted using a 5CV elution buffer (25mM Tris pH 8, 300mM NaCl, 500mM imidazole). Eluate was purified by size exclusion chromatography (SEC) on an AKTA PURE FPLC system, using either a Superose 6 Increase 10/300 GL column or a Superdex 200 Increase 10/300 GL column, with Tris-buffered saline (TBS; 25mM Tris pH 8, 150mM NaCl) at a speed of 0.75mL/min. Fractions corresponding to the peak trace were collected and combined for final purified protein. This was aliquoted into PCR tubes and stored at -80°C for future use.

### Lentivirus production, Transduction, & Cell line Creation

LV-TRE-VP64 human MyoD-T2A-dsRedExpress2 & PLX304-ACVRL1 lentivirus particles were prepared by first plating HEK293FT cells at 70% confluency on 0.1% gelatin coated 10 cm dish were transfected with 20 ug of shuttle vector, 15 ug of helper plasmids psPAX2, and 5 ug of pMD2.G using 120uL 1mg/mL PEI transfection reagent in Opti-MEM™. 16 hours later the media was replaced with 8 mL fresh HFF media. Virus containing cell culture supernatant was collected after 3 days and filtered through PES 0.45 um filters. Aliquots were used the day of collection, with extras stored at -80°C.

For generating HFF-iMyoD, HFFs were MyoD-T2A-dsRedExpress2 lentivirus supplemented with final concentration of 6 ug/mL of polybrene for two consecutive days. The next day media was replaced with HFF media with 0.5 ug/mL puromycin for selection. After two days, media was replaced with HFF media and allowed to recover till 80% confluency, where cells were passaged to 10 cm dishes for expansion and freezing.

For generating CHO-hALK1, CHO cells were grown to 70% confluence, were treated with the PLX304-ACVRL1 lentivirus supplemented with final concentration of 6 mg/mL of polybrene for two consecutive days. The next day media was replaced with CHO media with 5 ug/mL Blasticidin for selection, with media replaced every other day. After four days, media was replaced with CHO media and allowed to recover till 80% confluency, where cells were passaged to 10 cm dishes for expansion and freezing.

For generating WTC-11-MyoD, 0.7 to 1x10^6 cells of WTC-11 were transfected with 0.5µg AAVS1-TALEN-R plasmid, 0.5µg AAVS1-TALEN-L and 1µg of AAVS1-TRE3G-MyoD-T2A-dsRedExpress2 using Amaxa Lonza Human stem cell Kit #2. The cells were then plated with 5μM Rocki onto a 1:40 matrigel solution coated plate. Two days following the transfection, the cells were selected for Puromycin 0.5µg/ml for 2 days.

### Myogenic transdifferentiation assay for Screening designed proteins

HFF-MyoD cells were used to evaluate the effects of designed proteins on myogenic conversion. 96-well, flat-bottom, clear-bottom, black polystyrene tissue-culture-treated microplates were coated with 0.1% gelatin for at least 15 min at 37°C and 5% CO₂. Following removal of the gelatin solution, HFF-MyoD cells were seeded at 1 × 10³ cells per well in HFF medium and maintained at 37°C and 5% CO₂, with medium changes every other day.

At approximately 80% confluency (designated Day 0), the medium was replaced with serum-containing skeletal muscle medium (SMM+FBS) consisting of high-glucose DMEM supplemented with GlutaMAX^TM^, 10% FBS, 10% KnockOut Serum Replacement, 1% penicillin/streptomycin, and 1% non-essential amino acids (NEAA), together with 2 µg/mL doxycycline to induce MyoD expression. Cells were maintained in SMM+FBS with medium changes every other day. On Day 4, the medium was replaced with SMM without serum consisting of high-glucose DMEM supplemented with GlutaMAX, 20% KnockOut Serum Replacement, 1% penicillin/streptomycin, and 1% NEAA, containing 10 µM CHIR99021 and 100 nM of the indicated designed protein. Treatment medium was replaced every other day. On Day 8, cells were fixed and immunostained for desmin, myosin heavy chain (MHC), and ATP synthase, with nuclei counterstained using DAPI.

### Long-term myogenic maturation assay

For long-term maturation studies, cells were subjected to the same induction and treatment protocol described above, except that treatment with 100 nM designed protein in SMM−FBS containing 10 µM CHIR99021 and 10 µM SB-431542 was continued with medium replacement every other day for 30 days. At the endpoint, cells were fixed and immunostained for α-actinin, with nuclei counterstained using DAPI.

### Alk1 shRNA knockdown in Myogenic transdifferentiation assay

In the myogenic transdifferentiation assay Alk1 shRNA lentivirus (vectorbuilder) was transduced onto the reprogramming cells in during day 2 and 3 of 8 days myogenic transdifferentiation protocol at MOI of 10 without polybrene (replacing media each day).

### Myogenic iPSC differentiation assay

WTC-11-MyoD cells were seeded at 0.24 × 10 cells per well in 24-well, flat-bottom, clear-bottom, black polystyrene tissue culture-treated microplates coated with 1:20 Matrigel. Cells were maintained in mTeSR medium at 37°C with 5% CO₂, and the media was replaced daily. Upon reaching approximately 80% confluency (day 0), cells were switched to mTeSR supplemented with 2 µg/mL doxycycline to induce MyoD overexpression. On day 2, the medium was replaced with SMM supplemented with fetal bovine serum (SMM + FBS) and 2 µg/mL doxycycline, with subsequent medium changes performed every other day. On day 6, cells were switched to SMM without FBS (SMM − FBS) containing 100 nM of the respective designed proteins and 10 µM CHIR9902. SMM − FBS consisted of high-glucose DMEM supplemented with GlutaMAX™, 20% KnockOut™ Serum Replacement, 1% penicillin–streptomycin, and 1% NEAA. Treatment medium was replaced every other day while maintaining the respective protein and CHIR9902 concentrations. On day 10, cells were fixed and immunostained for Desmin, myosin heavy chain (MHC), and DAPI.

### Immunostaining & Fixation

Cells were washed with 1X phosphate buffered saline (1X PBS) 3 times before cells were fixed in 4% paraformaldehyde (diluted in PBS) for 15 min. Following fixation, cells were washed three times for five minutes with 1X PBS. Cells were then permeabilized in PBST (1X PBS + 0.05% Triton X-100) for 1 minute. After permeabilization coverslips were blocked in 1% BSA (1% BSA + 1X PBST) for at least 1 hour at RT. The appropriate primary antibody diluted in 1% BSA was then added to the cells and left overnight at 4°C. The primary antibody solution was then removed and cells were given three 5 min washes with PBST. Appropriate fluorescent secondary antibodies diluted (1:500) in 1% BSA were then added to cells for 45 minutes at RT. Cells were given three 5 min washes in PBST, then a 2 ug/mL solution of DAPI diluted in 1% BSA was added for 15 minutes at RT. Three final 5 min washes with PBST were given. Glycerol+n-propyl gallate mounting solution (90% Glycerol, 10% 10xPBS, 2% Propyl Gallate) was added either directly into wells for 96 well plates or added onto a glass slide for mounting coverslips. For 96 well plates, 96 well plate lid is parafilmed to prevent evaporation. For Glass coverslips, coverslips were put onto mounting solution, pressed down to without lateral movement, and edges of coverslips were sealed using clear nail polish. Stained cells were kept at 4°C until imaging was performed.

Primary Dilutions: Mouse Anti-Myosin Heavy Chain (MHC) (1:200), Mouse anti-ATP-B synthase (1:500), Rabbit Anti-Desmin (1:1000), Rat anti-a-Actinin (1:1000) Imaging of 96 well screening plates occurred at either a Leica or Nikon spinning disk microscope stitched (4x4) 20X images with 15% overlap, one z-plane, and auto focused using the Perfect Focus System (PFS) of each 96 well.

### Mito-Orange assay

Mito-Orange was diluted to 200 nM in the appropriate prewarmed (37°C) SMM-FBS media. Cells were rinsed once with prewarmed growth media then the Mito-Orange + SMM-FBS media was added and cells were incubated at 37°C and 5% CO2 for 60 minutes. Following treatment, cells were fixed and immunostained.

### Myogenic Transdifferentiation Assay analysis: Efficiency, Length, Multinucleation, Desmin intensity

All image analyses were performed using Fiji (ImageJ) with custom in-house macros developed for automated image processing and quantification.

### Conversion Efficiency

a Fiji macro was used to select and count the total number of areas above the intensity & 50 um^2^. The resulting mask was used to quantify the total number of nuclei in each image. Desmin-positive nuclei were then manually identified using the Multi-point/Counter tool and marked in each image. The number of marked nuclei was recorded as the total number of Desmin-positive nuclei. Conversion efficiency was calculated for each image as the percentage of Desmin-positive nuclei relative to the total number of nuclei.

### Desmin Intensity & Area

Several representative images were analysed benchmarking the threshold corresponding to the Desmin background in 96-well plate. A Fiji macro was then used to apply this threshold and generate a mask containing regions with Desmin intensity above background. The total Desmin-positive area and integrated fluorescence intensity were measured for each well and experimental condition. These measurements were normalized to the total number of nuclei in the corresponding image.

### Myotube Length

The length of Desmin-positive cells was measured manually in Fiji using the Freehand Line tool. For each Desmin-positive cell, a line was traced along the curvature of the cell from one end to the other, and the resulting length was recorded. Approximately 100 Desmin-positive cells were measured per image. For each image, the percentage of measured Desmin-positive cells with lengths above or below 300 µm was calculated.

### Myotube width

Width of desmin positive cells were determined utilizing the straight line tool in Fiji.

### Multinucleation

Each Desmin-positive myotube was identified and categorized according to the number of nuclei present as mononucleated, binucleated, containing 3-5 nuclei, containing 6-9 nuclei, or containing more than 10 nuclei using the Multi-point/Counter tool. All Desmin-positive myotubes within each image were scored. The resulting counts were used to determine the distribution of myotubes across the different multinucleation categories for each experimental condition.

### gp130mb & HFF & EA.hy926 Signaling

HFFs & EA.hy926 were grown to 80% confluency in a 24 well plate. Media was aspirated, than washed 3 times with PBS, then replaced with low glucose (1g/L) DMEM. 20 to 24 hours later, media was replaced with low glucose DMEM & pre-treatment with IL6 inhibitors (GP130mb or Ruxolitinib) was performed for 60 min in respective treatment condition wells. Then the media was replaced with treatment conditions (50 ng/mL IL-6, GP130mb, & Ruxolitinib) in low glucose DMEM for 15 min. Lysis then occurred as described in the “Sample Lysis” section.

### Sample Lysis & Western Blotting

For lysis, cells were washed with PBS once before adding freshly made complete western blot lysis buffer (900 uL WB lysis buffer stock, 270 uL 10% SDS, 15 uL Phosphatase inhibitor, 2 uL Dr. Nuclease) to each well (45 uL per well of 24-well plate & 180 uL per 35mm plate), scratched, then samples were placed into tubes each with 5% 2-Mercaptoethanol in 4x Laemmli buffer. For muscle cell lysis buffer differed (900 uL WB lysis buffer stock, 270 uL 10% SDS, 15 uL Phosphatase inhibitor, 10uL protease inhibitor, 6 uL Dr. Nuclease), scratched, then placed into tubes with 10% 2-Mercaptoethanol in 2X Laemmli buffer. Samples were boiled at 95°C for 8 min, then stored at -20°C.

WB Lysis buffer stock: 25 mM β-Glycerophosphate, 50 mM NaF, 10 mM Na Pyrophosphate, 1 mM Na3VO4, 1 mM PMSF Protease Inhibitor, 1 Pierce Protease Inhibitor mini Tablet per 10 mL, all dissolved in Zhan’s buffer (20 mM Tris-HCl, 30 mM NaCl, 15% Glycerol, 1% Triton X-100 in ddH2O).

The following day, Lysates were boiled at 95°C for 7 minutes. After boiling, samples were vortexed for 5 seconds and spun down. For 15 well gels, 10 uL of lysate per sample was loaded per well and for 10 well gels 15 uL of sample was loaded. Gels were run for 40 minutes at 200V in a running buffer (1.44% Glycine, 0.3% Trizma, 0.1% SDS, 98.16% deionized water) using a Bio-Rad PowerPac Basic electrophoresis power supply. Following electrophoresis, gels were soaked in a transfer buffer (20% methanol, 0.24% Tris-HCl, 0.8% NaCl, 0.1% Tween-20, in deionized water) and transferred onto a nitrocellulose membrane for 12 minutes at 25V using a Bio-Rad TransBlot Turbo system. Following transfer, the membrane was blocked in 5% BSA in 1X TBST (0.242% Tris-HCL, 0.8% NaCL, 0.1% TWEEN 20 in deionized water at pH 7.4) for 1 hour at RT on a shaker, then incubated in primary antibodies overnight at 4°C on a rocker. Primaries were then recovered to be reused and three 5-minute washes with 1X TBST were performed before adding HRP-conjugated secondary antibodies (1:5000 dilution, except for S6 & H3 at 1:10000) in 5% BSA in 1X TBST for 1 hour at RT on a shaker. Three more 5-minute washes in 1X TBST were then performed on a shaker. A 50/50 solution of hydrogen peroxide and HRP substrate was added to membranes immediately before imaging on a Bio-Rad ChemiDoc MP Imaging System. Western Blot images were analyzed using Fiji (ImageJ).

Primary Dilutions: 1:5000: Rabbit Anti-S6, Rabbit Anti-Histone H3. 1:1000: Rabbit Anti-b-actin, Rat anti-a-Actinin, Rabbit anti-Myogenin, Goat Anti-Human ALK1, Rabbit Anti-ETV1, Rabbit Anti-ETV4, Rabbit Anti-HMGA2, Rabbit Anti-HA-Tag. 1:500: Mouse Anti-MHC, Rabbit anti-Phospho-SMAD1/5 (Ser463/465).

### Single nuclei RNA sequencing

Single-cell combinatorial-indexing RNA-sequencing (sci-RNA-seq) protocol is described previously (Cao et al., 2019; Martin et al., 2023). Briefly, sci-RNA-seq relies on thawed nuclei permeabilized with 0.2% Triton X-100 in nuclei wash buffer (0.333M sucrose, 0.1% Triton X-100, 3 mM MgCl2, 10% 10x DPBS, dissolved in nuclease-free water) for 3 min on ice, and briefly sonicated to reduce nuclei clumping. Nuclei were then distributed across 96-well plates. A first molecular index is introduced to the mRNA of the nuclei within each well, with in situ reverse transcription (RT) incorporating the unique molecular identifiers (UMIs). Then all nuclei were pooled and redistributed to multiple 96-well plates in limiting numbers (e.g., 10 to 100 per well) and a second molecular index was introduced by hairpin ligation. Second strand synthesis, tagmentation, purification and indexed PCR; Library purification and sequencing is performed. All libraries were sequenced on an Illumina NovaSeq 6000 system. Base calls, downstream sequence processing and single-cell digital-expression matrix generation steps were similar as described in the sci-RNA-seq3 paper (Cao et al., 2019), which is STAR v.2.5.2b aligner used with default settings and gene annotations (GRCh38-primary-assembly, gencode.v27). Uniquely mapping reads were extracted, and duplicates were removed using the UMI sequence, reverse transcription index, hairpin ligation adaptor index and read 2 end-coordinate (that is, reads with identical UMI, reverse transcription index, ligation adaptor index and tagmentation site were considered duplicates).

Sci-RNA-seq Data Analysis: All low-quality reads were removed from the data by setting UMI cutoff to greater than 200 and removing all mitochondrial reads. Following the Monocle3 workflow, data underwent normalization by size factor, preprocessing, dimension reduction (UMAP algorithm), and unsupervised graph-based clustering analysis (Leiden Algorithm). Certain clusters from the initial analysis were selected for further sub-clustering, and the previous analysis repeated. Pseudotime analysis also was done with Monocle3 following the default workflow, which included learning the graph, ordering the cells, and plotting the trajectory over UMAP.

### Barrier Score

HFF-MyoD were plated and differentiated just as in the Myogenic transdifferentiation assay, SMM+FBS supplemented with 2 ug/ml doxycycline occurred for all days. Cells were harvested and nuclei were isolated on day 0 (8,616 nuclei), day 7 (4,643 nuclei), & day 14 (860 nuclei), followed by single nuclei Sci-sequencing. Monocle3 preprocessing and dimensionality reduction methods were used to generate UMAPs. UMAP identified three major clusters of cells: Resting Fibroblast (Cluster 1), Myofibroblast / Activated fibroblast (Cluster 2), and Myogenic State (Cluster 3). Important identifier genes for each cluster were: Resting Fibroblast (Cluster 1) {FGF12, TCF21 , PDGFRA , DCN , LUM}; Myofibroblast / Activated fibroblast (Cluster 2) {COL8A1 (collagen type VIII alpha 1), contractile myofibroblast markers (ACTA2 / -SMA, TAGLN / SM22), and matrix remodeling factors (POSTN , FN1 , MMP11 , THBS1)}; Myogenic State (Cluster 3) {TNNT2 (cardiac/skeletal troponin T2), ACTC1 (alpha actin), MYOD1 , MYOG (myogenin), DES (desmin), and TTN (titin)}. To quantify the molecular resistance and refractory state hindering direct myogenic reprogramming at single-cell resolution, a composite Barrier Score was computed using Monocle 3. The barrier signature was constructed from a curated panel of candidate barrier and transdifferentiation-inhibitory genes, including master epithelial-to-mesenchymal transition (EMT) and fibrogenic transcription factors (*TWIST1*, *TWIST2*, *ZEB1*, *SNAI2*), extracellular matrix components (*COL1A1*, *FN1*), non-canonical Wnt signaling (*WNT5A*), and myogenic inhibitors (*ID1*). For each single nuclei , the barrier module score was calculated as the mean normalized expression across all detected barrier genes:

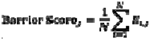

where represents the normalized log-transformed expression level of barrier gene in nuclei. The resulting continuous metric was assigned to the cell metadata (colData) to project barrier intensity across the UMAP manifold, track barrier dynamics along the pseudotime differentiation trajectory, and serve as the reference vector for genome-wide Pearson correlation screening to identify upstream transcription factors and regulatory drivers sustaining the reprogramming bottleneck.

### Inflammatory score

To assess cellular inflammatory activation at single-cell resolution, an inflammatory gene signature score was calculated using the AddModuleScore function in the Seurat R package (v4.3.0 / v5.0). The signature comprised a curated panel of 266 canonical inflammatory genes covering interleukins (IL family), chemokines (CXCL and CCL families), tumor necrosis factor superfamily members (TNF , TNFRSF ), interferons and interferon-stimulated genes (IFN , IFI , IFIT), NF-κB pathway components (NFKB1 , NFKB2 , NFKBIA), inflammasome effectors (NLRP family), Toll-like receptors (TLR family), and key mediators of inflammation (C3 , PTGS2 , STAP2 , and CRP) (**Table EV11**). Briefly, for each individual cell, the module score was computed by calculating the average expression of the target inflammatory gene set subtracted by the aggregated expression of randomly selected control gene sets. To prevent bias related to gene expression magnitude, control genes were binned into 24 expression bins based on their average expression across the entire transcriptome. The resulting normalized module score quantified cell-specific inflammatory enrichment relative to baseline and was compared across experimental conditions ( Fibroblast , No MB , and C6 Cocktail ), with distributions visualized using projection (UMAP) feature plots. Number of single nuclei analysed for the inflammatory score per condition: fibroblast_control (Total:12,132), No MB (Total: 31,878), C6 Cocktail (Total: 46,791).

### Alk1-IGF2_Endotag4 mediated Alk1 degradation

1. For western blot or immunostaining analysis, CHO-ALK1 cells were plated on a full 24 well plate (equivalent seeding density for standard maintenance) in CHO media. For immunofluorescence analysis, cells were plated on glass coverslips that were coated with 0.1% gelatin (double seeding density as for standard maintenance.
2. At 70% confluence media was changed for fresh CHO media and treatment with 100nM ALK1mb-IGF2_Endotag4 or IGF2_Endotag4_ALK1mb was added to applicable wells.
3. 24 hours later, for non-washout conditions (including PBS) coverslips were fixed while normal wells were lysed for western blot. Washout conditions, cells were washed 3 times with PBS before fresh CHO media was added.
4. 24 hours later rest of coverslips were fixed while normal wells were lysed for western blot.
5. Lysates were then run in western blot, and coverslips were immunostained for ALK1.
6. Immunostained coverslips were analyzed in Fiji (ImageJ) by taking the total integrated ALK1 intensity of the images, which was then normalized for the number of nuclei.

### Seahorse assay

HFF-MyoD cells were seeded into a 0.1% gelatin coated 96-well Seahorse XFe96 culture plate at a density of 2.0 X 10^4 cells per well. Cells were grown in HFF maintenance media until they reached 80% confluency, at which point the transdifferentiation with the same procedure as with the screening assay up to day 8. Two hours prior to the seahorse assay, cells were rinsed with PBS and given fresh SMM-FBS media. Cells were then rinsed twice with prewarmed seahorse assay media (XF DMEM, 2mM L-Glutamine, 10mM Glucose, 2mM Sodium Pyruvate) and incubated for 30 minutes in 200 uL assay media per well at 37°C (No CO2) in the XF Prep Station. After incubation cells were washed one final time, and 180 uL of warmed assay media was added. An XFe96/XF Pro Cell sensor cartridge was loaded with three treatments, 20 uL of 2.5 uM oligomycin A in port A, 22 uL of 1 uM Carbonyl cyanide 4-(trifluoromethoxy)phenylhydrazone (FCCP) in port B, and 25 uL of 2.5 uM Antimycin A + 2.5 uM Rotenone in port C. All drugs were dissolved in seahorse assay media. Port D was filled with 28 uL of assay media as required by the manufacturer. The assay was run at 37°C in an Agilent Seahorse XFe96 Analyzer with four measurements for each treatment were taken with 3 minutes of mixing, zero seconds of waiting, and 3 minutes of measuring. After the assay, the media was removed from the cell culture plate and it was frozen overnight at -20°C. The next day cells were thawed for 20 minutes, 40 uL of 0.01% SDS was added to each well and the plate was frozen at -80°C for 30 minutes. Cells were then thawed for 15 minutes and 40 uL of Hoechst solution (6 ml 1M NaCl, 60 ul 0.1 M EDTA, 60 ul 1M Tris HCl, 2.4ul 20 mM Hoechst solution) was added to each each well, and the plate was left to incubate for 1 hour at 37°C (No CO2). A Wallac EnVision 2104 Multilabel Reader analyzer was used to take three separate measurements of the Hoescht intensity in each well. These values were then averaged and the empty corner well values subtracted, then used for normalizing the measured oxygen consumption rate values for each well. The data was analyzed using Agilent Seahorse Analytics software.

### Bulk RNA sequencing

Cells were grown as per the described conditions, making sure that there is at least 80% confluence before doing Bulk RNA sequencing lysis & that all cells are ready for lysis on the same day. Bulk sequencing was performed as per Plasmidsaurus, or BGI. Briefly, cells were lysed in DNA/RNA Shield for Plasmidsaurus while extracted RNA was used for BGI. Differential gene analysis was performed using Partek.

### RNA-seq differential gene expression and volcano plot analysis

RNA-sequencing data were analyzed using Partek Flow v11.0.24.0624. Raw sequencing reads were aligned to the human reference genome (hg19) using STAR v2.7.8a with default parameters. Aligned reads were quantified using the Quantify to Annotation Model function against the Ensembl release 100 hg19 annotation model (hg19_ensembl_release100_v2). Features with fewer than 10 reads summed across all samples were excluded from downstream analysis. Gene-level counts were normalized using the DESeq2 median-ratio method, and differential gene expression between no mb, C2-DPC, C6-DPC versus fibroblast and C6-DPC-treated versus no-minibinder samples was determined using DESeq2 in Partek Flow. DESeq2 modeled gene counts using a negative binomial distribution, and P-values were calculated for differential expression between the two conditions. GSE352439 data was used for volcano plot analysis.

Differential expression results, including gene identifiers, P-values, and log2 fold changes, were exported from Partek Flow and further analyzed in R. Volcano plots were generated using the ggplot2, dplyr, and ggrepel packages. Genes were classified as upregulated when log2 fold change > 1 and P < 0.05 and downregulated when log2 fold change < −1 and P < 0.05, corresponding to an absolute fold change greater than two. Genes not meeting both thresholds were considered nonsignificant for visualization. P-values equal to zero were replaced with 1 10 before −log10 transformation. The volcano plot displays log2 fold change on the x-axis and −log10(P-value) on the y-axis, with dashed lines indicating the thresholds of log2 fold change = ±1 and P = 0.05. Selected genes of biological interest were annotated using ggrepel.

### Promoter sequence extraction and transcription factor motif enrichment analysis

Promoter motif analysis was performed on differentially upregulated genes during C6-DPC– mediated myogenic conversion (the GEO accession number: GSE352439). Duplicate gene symbols were removed, and genomic annotations, including transcript transcription start sites (TSSs), chromosome assignments, and strand information, were retrieved from Ensembl for the human GRCh38 reference genome using the R package biomaRt. Annotations were restricted to chromosomes 1–22, X, and Y. For each gene–chromosome–strand group, the most upstream annotated TSS was selected, corresponding to the minimum genomic coordinate on the positive strand and the maximum coordinate on the negative strand. Strand-aware promoter regions comprising 2,000 bp upstream and 200 bp downstream of the selected TSS were defined using GenomicRanges. Promoter sequences were extracted from the hg38 reference genome using BSgenome.Hsapiens.UCSC.hg38 and exported in FASTA format.

Human transcription factor motifs were retrieved from JASPAR2020 (Fornes et. al., 2020) using TFBSTools, retaining the latest available version of each motif. Position frequency matrices were converted to position weight matrices, and promoter sequences were scanned using motifmatchr. A background set containing one synthetic sequence per promoter was generated with matched sequence lengths and fixed nucleotide probabilities of 0.29 for A and T and 0.21 for C and G, corresponding to an expected GC content of 42%. Random sequence generation used a seed of 123. Motif scanning was performed separately for promoter and background sequences using subject-derived nucleotide frequencies.

For each motif, sequences were classified according to the presence or absence of at least one predicted binding site. Motif enrichment was assessed using a one-sided Fisher’s exact test comparing motif-positive and motif-negative sequences between the promoter and background sets. P values were adjusted for multiple testing using the Benjamini–Hochberg procedure, with an adjusted P value below 0.05 defining statistical significance. Total 174 motifs are significant with padj< 0.05, The 25 most significant motifs were selected for visualization of log₂-transformed odds ratios, with transcription factor family annotations.

### Plasmid Cloning

PLX304-ACVRL1 was generated by gateway cloning using the Gateway™ LR Clonase™ II Enzyme mix kit with the pDONR223-ACVRL1 Entry Clone and PLX304 Destination vector. Cloning was performed as per the Gateway™ LR Clonase™ II Enzyme mix kit’s procedure. AAVS1-TRE3G-MyoD-T2A-dsRedExpress2 was generated by first PCR amplifying the MyoD-T2A-dsRedExpress2 fragment region of the LV-TRE-VP64 human MyoD-T2A-dsRedExpress2 with inserted MluI and AccI restriction sites using the MyoD-T2A-dsRedExpress2 Gibson Assembly Forward and Reverse Primers. Using the MluI-HF & AccI restriction enzymes the AAVS1-TRE3G-EGFP was digested. The AAVS1-TRE3G fragment was purified by Qiagen DNA Gel Extraction. Then the two fragments were Gibson assembled using the Gibson Assembly® Master Mix as per New England Biolabs “Gibson Assembly® Master Mix Assembly Protocol (NEB #E2611)”. The finished reaction mixture was transformed into DH5a cells for plasmid isolation and purification.

## 3-D Force experiments

### Differentiation, purification, and expansion of iPSC-derived skeletal muscle myoblasts

iPSC-derived myoblasts (iPSC-MBs) were generated from the UC3-4 DMD null and isogenic control UC3-4 iPSC lines (Smith et al., 2022), using a multi-stage protocol adapted from previously published protocol (Chal et al., 2015). From Day 0 to Day 7, cultures were fed daily with 2 mL of fresh medium. From Day 7 onwards, the media was refreshed every 48 hours. Basal DMEM/F12 media: DMEM/F-12, supplemented with 1x non-essential amino acids.

1. Initial expansion of induced pluripotent stem cells (iPSCs) was performed in mTeSR Plus on 6-well plates (seeded at 15,000 cells/cm²) until cultures achieved ∼40% confluence.
2. Day 0: Differentiation was initiated by exchanging the media for Basal DMEM/F12 media supplemented with and 1x Insulin-Transferrin-Selenium (ITS), 3 µM CHIR99021 and 0.2 µg/mL LDN193189.
3. Day 2: Media switched to Basal DMEM/F12 media supplemented with 1xITS, 3 µM CHIR99021, 0.2 µg/mL LDN193189 & 20 ng/mL bFGF.
4. Day 5: Media switched to Basal DMEM/F12 media supplemented with 15% KnockOut™ Serum Replacement, 0.2 µg/mL LDN193189, 20 ng/mL bFGF, 2 ng/mL IGF-1, and 10 ng/mL HGF.
5. Day 7: Media switched to Basal DMEM/F12 media supplemented with 15% KnockOut™ Serum Replacement, 2 ng/mL IGF-1, and 10 ng/mL HGF.
6. Day 28: To expand the myogenic population, a single well was mechanically dissociated into aggregates and replated into a 225^2 cm cell culture flask, coated with 1:60 matrigel. Expansion was carried out in Skeletal Muscle Growth Media (SKGM: DMEM with high glucose & GlutaMAX™ Supplement, 10% FBS, 100 nM Insulin, 40 ng/mL bFGF2, and 1 µM Dexamethasone).
7. Day 32: single-cell suspensions were generated by filtration (40 µm) and sorted via fluorescence-activated cell sorting (FACS) to isolate Nerve Growth Factor Receptor-positive (NGFR+) progenitors. Experiments utilized cells from passages 4–5.

### Generation of 3D engineered muscle tissues (EMTs)

Engineered muscle tissues (EMTs) were generated using the Mantarray platform (Curi Bio) following an adapted fabrication protocol.

1. Day -1: A co-culture cell pellet was prepared by combining iPSC-derived myoblasts and human dermal fibroblasts at a 9:1 ratio. Cells were resuspended to achieve a final density of 420,000 cells per 60 µL hydrogel mixture composed of 1.3 mg/mL rat collagen I, 20% Matrigel, 10% Minimum Essential Medium (MEM), and 10% DMEM with high glucose & GlutaMAX™ Supplement.This cell-hydrogel suspension was pipetted directly into the Mantarray casting trenches and polymerized for 60 minutes at 37 °C and 5% CO2.
2. Following polymerization, 1 mL of SKGM was added per well for 24-hours.
3. Day 0: Constructs were then transferred to fresh 24-well plates containing 1.5 mL of differentiation medium (DMEM with high glucose & GlutaMAX™ Supplement, and 2% Horse Serum. For treatment groups, this medium was supplemented with 100 nM C6 cocktail.
4. Maintenance involved replacing half the media volume (± C6 cocktail) every 2–3 days.

### Measurement of EMT contraction

Contractile function was assessed using the Mantarray platform (Curi Bio). The 24-well EMT plate was loaded into the instrument’s measurement chamber, maintained at 37°C, and a stimulation lid assembly was inserted to position electrodes within each well, enabling simultaneous electrical pacing of all tissues. Magnetic field flux was recorded at a sampling rate of 100 Hz and converted to post deflection (mm) using the manufacturer’s proprietary magnet localization algorithm. These deflection data were subsequently converted to contractile force to generate force-time traces for analysis. Tissues underwent three distinct stimulation protocols utilizing biphasic electrical pulses (5 ms duration, 100 mA). All raw force data were processed using the Engineered Muscle Contractility Analysis Pipeline (EMCAP) version 1.0, a custom open-source Python based peak-finding and analysis package (Barrett, 2026). Twitch force and kinetics were evaluated using a Twitch Protocol (0.2 Hz), consisting of 10 individual pulses separated by 5-second rest intervals; active twitch force was calculated as the average of these ten contractions. Tetanic force and kinetics were determined using a Force-Frequency Protocol, where tissues were subjected to 2-second stimulation trains at increasing frequencies (1, 2, 3, 5, 10, 15, 20, 30, 40, 60, 80, and 100 Hz), with a 16-second rest period between each step.

### EMT Immunohistochemistry and immunocytochemistry

1. To determine the effective physiological cross-sectional area (CSA) for specific force calculations and to assess myotube morphology, EMTs were fixed on Day 14 in 4% paraformaldehyde (24 h, 4°C).

1. Tissues were cryoprotected in 20% sucrose (w/w) for 24 hours, embedded in Optimal Cutting Temperature (OCT) compound, and stored at -80°C.
2. Transverse cryosections (30 µm) were blocked for 1 hour at room temperature in blocking buffer (2% BSA, 5% goat serum, 0.25% Triton X-100).
3. Sections were incubated overnight at 4°C with primary antibodies against MHC (1 µg/mL) and Dystrophin (2 µg/mL).
4. Following primary incubation, sections were stained Goat anti-mouse 488, Goat anti-rabbit 546, Phalloidin 647, and Hoechst (overnight, 4°C) and imaged using a Yokogawa W1 spinning disk confocal microscope equipped with a 20x air objective.
5. Image processing was automated using custom Fiji (ImageJ) macros. Image files were imported and aligned using the SIFT (Scale-Invariant Feature Transform) algorithm to correct for channel drift. To accurately segment individual myotubes, a "fusion" image was generated by averaging the background-subtracted MF-20 and Phalloidin signals. This fusion strategy was critical for maintaining segmentation fidelity. These fused images were segmented using StarDist plugin, a deep-learning-based object detection tool (Weigert & Schmidt, 2022). Individual and sum of myotube CSA was calculated from these segmented regions and used to normalize active force to specific force (mN/mm²).

### Statistical analysis

p-value significance tests were determined using the two-sided Student’s t-test.

Motif enrichment was assessed using a one-sided Fisher’s exact test comparing motif-positive and motif-negative sequences between the promoter and background sets. P values were adjusted for multiple testing using the Benjamini–Hochberg procedure, with an adjusted P value below 0.05 defining statistical significance.

## Data availability

All scRNAseq data have been submitted to the Harvard Dataverse at, https://doi.org/10.7910/DVN/2B5OGU. Bulk RNA-seq Data have been submitted to NCBI Gene Expression Omnibus (GEO) under the GEO accession number-GSE330553 and GSE352439.

## Acknowledgements

We thank all members of the HRB lab for valuable discussions. We thank Isabella Buenaflor and Kevin Huang and Samriddhi Jhajharia of Ruohola-Baker lab for experimental assistance. We also thank Ria Aggarwal of the Mack lab for assistance with the 3D force experiments, Dr. Mary Regier of the ISCRM Genomics Core for genomics support, and Dr. Dale Hailey and the Garvey Imaging Core for imaging support.

**Extended Figure 1** (A) A schematic showing the recombinant MyoD transcription factor used in our study with N -ter VP64 transactivation domain, HA tag and on c-term linked to dsRED with a cleavable T2A site. Western blot analysis shows that HEK cells transfected with MyoD plasmid treated with doxycycline show expression of MyoD protein, Beta actin is used as a housekeeping gene. (B) A bar graph showing the Myod/no of nuclei intensity in HFF-iMYOD cells immunostained for MyoD antibody and DAPI.

(C) tet-on inducible MyoD Human Foreskin FIbroblast were treated with Doxycycline for 7 days and 14 days, to reprogram into muscle cells, where single cell nuclei sequencing was performed. (D) UMAP showing all single nuclei RNA sequenced at day0, day7, day14.

(E) UMAP showing single nuclei RNA sequencing of all treatments excluding cluster 5. Cluster 5 is a separate cluster of cells from fibroblast HFFiMYOD uninduced cells (cluster1). (F) UMAP showing D0 (resting fibroblast), D7 (tKM) and d14 (tSKM) cells.

(G)Pseudotime analysis using Monocle3. (H) A dot plot showing resting, activated fibroblast, and muscle markers in the three specific clusters. (I) A schematic showing the antimyogenic or barriers of myogenic gene expression was scored to calculate the barrier score of myofibroblast/ activated fibroblast versus the myogenic cluster. The bar graph shows regulatory drivers of anti-myogenic (barrier score) of myofibroblast/ activated fibroblast versus the myogenic cluster.

(J) A schematic showing different time points [d4,d6,d8,d10,d14] of Doxycyline (dox) treatment followed by release in the media lacking serum and dox treatment [d0,d6,d8,d10,d12]. (K) A bar graph showing the various time point treatment and conversion efficiency (% of Desmin positive nuclei) was calculated from cells fixed and immunostained for the Desmin and Myosin heavy chain and DAPI.

(L) Western blot analysis of day 0 and day 8 tSKM showing muscle marker protein expression level of alpha-actinin, Myosin heavy chain(MHC) and Myogenin and two housekeeping genes Histone 3 and S6.(M) Conversion efficiency of muscle reprogramming in shRNA knockdown of ALK1 in transdifferentiation.

**Extended Figure 2** (A) A schematic showing fibroblast to tSKM conversion in presence of various combinations of design protein cocktail components treatment.(B) A bar graph showing myotube area calculated using imageJ from cells stained for Desmin and DAPI where the Desmin area over the same no of nuclei was calculated under different design minibinders treatments in d8 tSKM. (C)A bar graph showing conversion efficiency calculated from cells stained for Desmin and DAPI under different design minibinders treatments in d8 tSKM. (D) Immunofluorescence images of under no mb (control) and C2-DPC and C6-DPC treated d8 tSKM immunostained for the Desmin (green), MHC( red) and DAPI (DAPI) in 96 well where 4 by 4 stitched images are shown.(E) Immunofluorescence images of no mb (control) and C6-DPC treated d8 tSKM immunostained for the Desmin (green), MHC( red) and DAPI (DAPI) showing multinucleated myotubes. Scale bar=200um and 20um

**Extended Figure 3** (A)Volcano map showing skeletal muscle gene expression upregulated in no mb, C2-DPC and C6-DPC 8day tSKM over the uninduced HFF iMYOD fibroblast. (B) Unique and common (red) gene expression found in C2-DPC and C6-DPC 8day tSKM over the fibroblast. (C) Density of nuclei in single nuclei sequencing of fibroblast, no mb and C6-DPC treated mononucleated cells. (D) A volcano map showing pseudobulk data where upregulated and downregulated genes are represented in C6-DPC versus no mb mononucleated cells upon 8 days of transdifferentiation (E) Schematic showing C6-DPC alone, and C6-DPC+TNF-a treatment during 8 days of reprogramming. A bar graph showing C6-DPC+TNF-a decreases the conversion efficiency compared to the C6-DPC alone treatment. (F) Western blot analysis shows gp130mb downregulates the IL6 mediated pSTAT3 level, bar graph shows the quantification of normalised pSTAT3 level from western quantification. (G) Western blot analysis shows Ruxolitinib downregulates the IL6 mediated pSTAT3 level, bar graph shows the quantification of normalised pSTAT3 level from western quantification.

**Extended Figure 4** (A) KEGG analysis of genes upregulated in C6-DPC versus no minibinder.

(B) Biological Process GO terms, analysis of genes upregulated in C6-DPC versus no minibinder. (C) Reactome analysis of genes upregulated in C6-DPC versus no minibinder. (D) qPCR mRNA expression of genes in C6-DPC over no mb tSKM. (E) Top transcription factors based on their score from ENCODE ChIP-seq library in upregulated gene expression in C6-DPC.

**Extended Tables EV1:** Minibinder sequences used in the primary screen (Fig 1F)

**Extended Tables EV2:** Desmin intensity of the primary screening of minibinders combination (Fig 1H)

**Extended Tables EV3:** Differential gene analysis of nomb, C2-DPC, and C6-DPC versus HFF-iMyoD (Fig EV3A)

**Extended Tables EV4:** Heat map in the fig 3H, analysis of the indicated gene signatures

**Extended Tables EV5:** Sequences of minibinders targeting inflammatory pathways (Fig 4E)

**Extended Tables EV6:** Desmin intensity of C6-DPC in combination with inflammatory pathways minibinders (Fig 4F)

**Extended Tables EV7:** Differentially expressed genes, C6-DPC versus nomb (Fig 6B)

**Extended Tables EV8:** Transcription factor (TF) motif enrichment analysis in promoters of C6-DPC upregulated genes (Fig 6C)

**Extended Tables EV9:** ETV1 TF binding sites predicted in C6-DPC upregulated genes targets

**Extended Tables EV10:** ETV4 TF binding sites predicted in C6-DPC upregulated genes targets

**Extended Tables EV11:** List of Genes used in calculating inflammatory score

