## Supplementary material for "Designed Minibinders Rewire Receptor Signaling to Enable Functional Human Myogenic Reprogramming": EV

### Figure-EV1

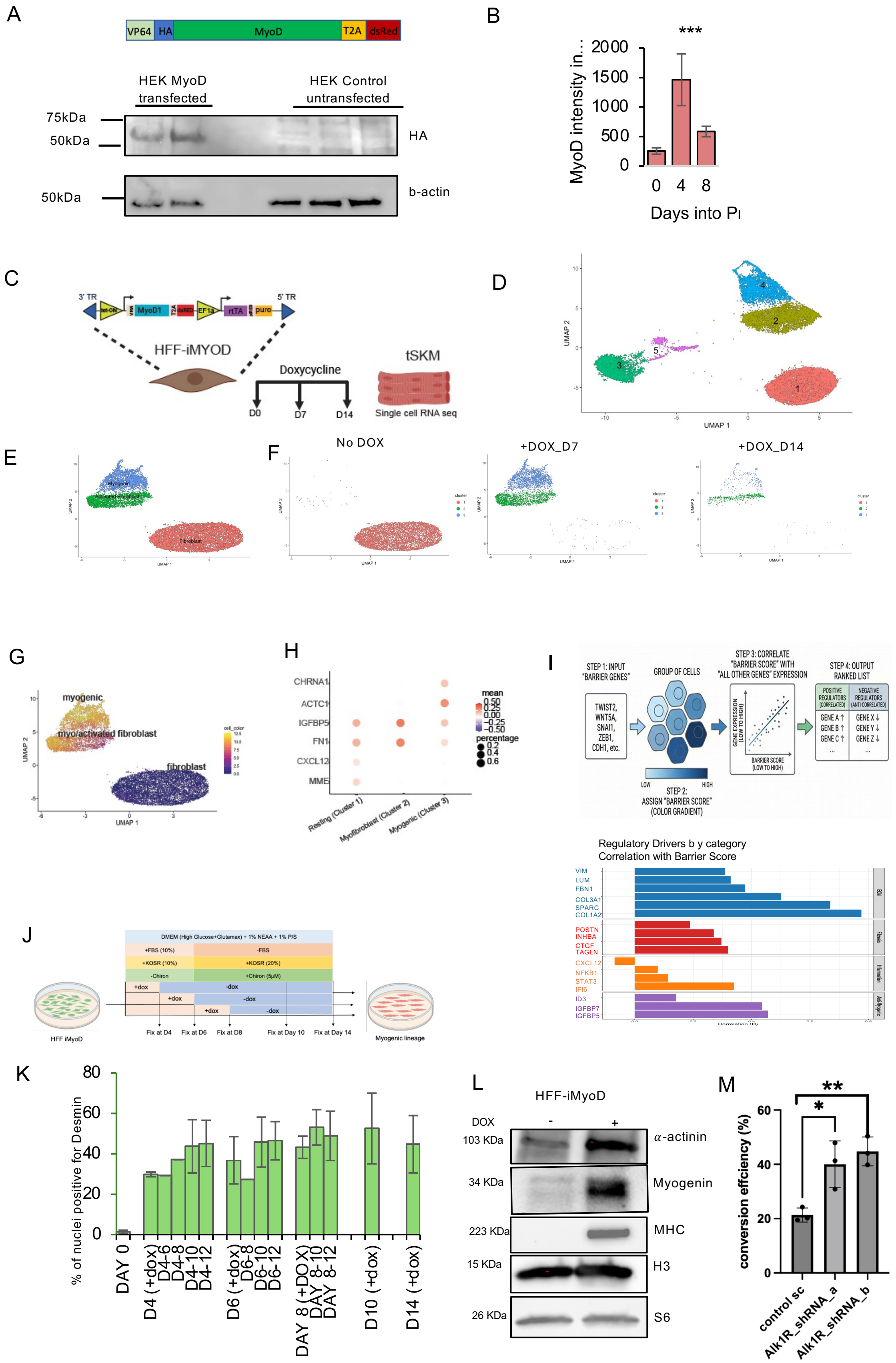

A
VP64
HA
MyoD
T2A
dsRed
HEK MyoD transfected
HEK Control untransfected
75kDa
50kDa
HA
50kDa
b-actin
B
***
MyoD intensity in...
Days into PI
C
3' TR
5' TR
HFF-iMYOD
Doxycycline
D0
D7
D14
tSKM
Single cell RNA seq
D
UMAP 2
UMAP 1
E
F
No DOX
+DOX_D7
+DOX_D14
G
myogenic
myo/activated fibroblast
fibroblast
H
CHRNA1
ACTC1
IGFBP5
FN1
CXCL12
MME
mean
percentage
Resting (Cluster 1)
Myofibroblast (Cluster 2)
Myogenic (Cluster 3)
I
STEP 1: INPUT "BARRIER GENES"
TWIST2, WNT5A, SNAI1, ZEB1, CDH1, etc.
GROUP OF CELLS
STEP 2: ASSIGN "BARRIER SCORE" (COLOR GRADIENT)
STEP 3: CORRELATE "BARRIER SCORE" WITH "ALL OTHER GENES" EXPRESSION
STEP 4: OUTPUT RANKED LIST
POSITIVE REGULATORS (CORRELATED)
NEGATIVE REGULATORS (ANTI-CORRELATED)
Regulatory Drivers b y category
Correlation with Barrier Score
VIM
LUM
FBN1
COL3A1
SPARC
COL1A2
POSTN
INHBA
CTGF
TAGLN
CXCL12
NFKB1
STAT3
IFI6
ID3
IGFBP7
IGFBP5
J
DMEM (High Glucose+Glutamax) + 1% NEAA + 1% P/S
+FBS (10%)
-FBS
+KOSR (10%)
+KOSR (20%)
-Chiron
+Chiron (5μM)
+dox
-dox
Fix at D4
Fix at D6
Fix at D8
Fix at Day 10
Fix at Day 14
HFF iMyoD
Myogenic lineage
K
% of nuclei positive for Desmin
DAY 0
D4 (+dox)
D4-6
D4-8
D4-10
D4-12
D6 (+dox)
D6-8
D6-10
D6-12
DAY 8 (+DOX)
DAY 8-10
DAY 8-12
D10 (+dox)
D14 (+dox)
L
HFF-iMyoD
DOX
-
+
103 KDa
α-actinin
34 KDa
Myogenin
223 KDa
MHC
15 KDa
H3
26 KDa
S6
M
conversion efficiency (%)
**
*
control sc
Alk1R_shRNA_a
Alk1R_shRNA_b

### Figure-EV2

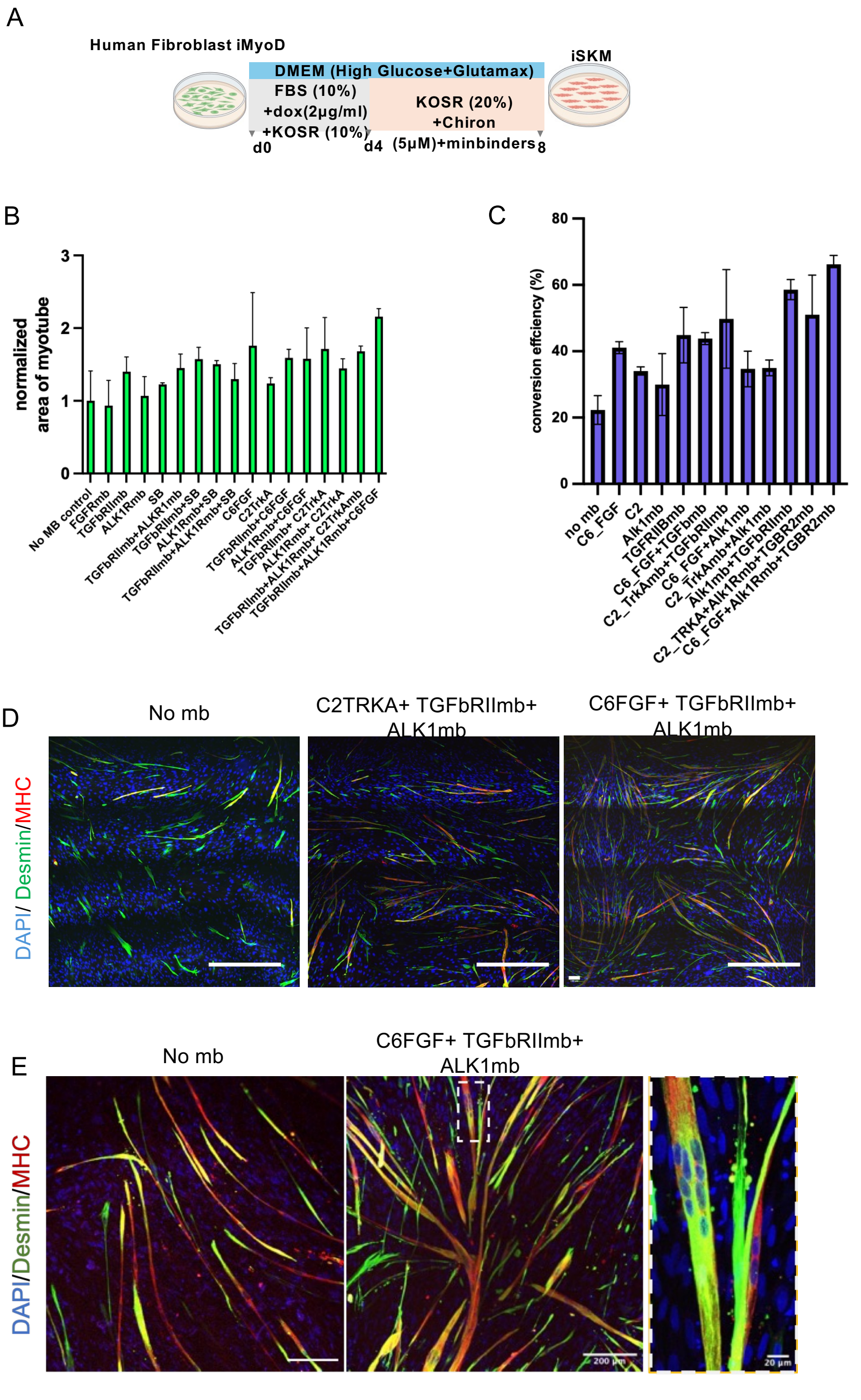

A
Human Fibroblast iMyoD
DMEM (High Glucose+Glutamax)
FBS (10%)
+dox(2µg/ml)
+KOSR (10%)
KOSR (20%)
+Chiron
(5µM)+minbinders
iSKM
d0
d4
8
B
normalized area of myotube
No MB control
FGFRmb
TGFbRIImb
ALK1Rmb
SB
TGFbRIImb+ALKR1mb
TGFbRIImb+SB
ALK1Rmb+SB
TGFbRIImb+ALK1Rmb+SB
C6FGF
C2TrkA
TGFbRIImb+C6FGF
ALK1Rmb+C6FGF
TGFbRIImb+ C2TrkA
ALK1Rmb+ C2TrkA
TGFbRIImb+ALK1Rmb+ C2TrkAmb
TGFbRIImb+ALK1Rmb+C6FGF
C
conversion effciency (%)
no mb
C6_FGF
C2
Alk1mb
TGFRIIBmb
C6_FGF+TGFbmb
C2_TrkAmb+TGFbRIImb
C6_FGF+Alk1mb
C2_TrkAmb+Alk1mb
Alk1mb+TGFbRIImb
C2_TRKA+Alk1Rmb+TGBR2mb
C6_FGF+Alk1Rmb+TGBR2mb
D
No mb
C2TRKA+ TGFbRIImb+ ALK1mb
C6FGF+ TGFbRIImb+ ALK1mb
DAPI/ Desmin/MHC
E
No mb
C6FGF+ TGFbRIImb+ ALK1mb
DAPI/Desmin/MHC
200 µm
20 µm

Figure-EV3

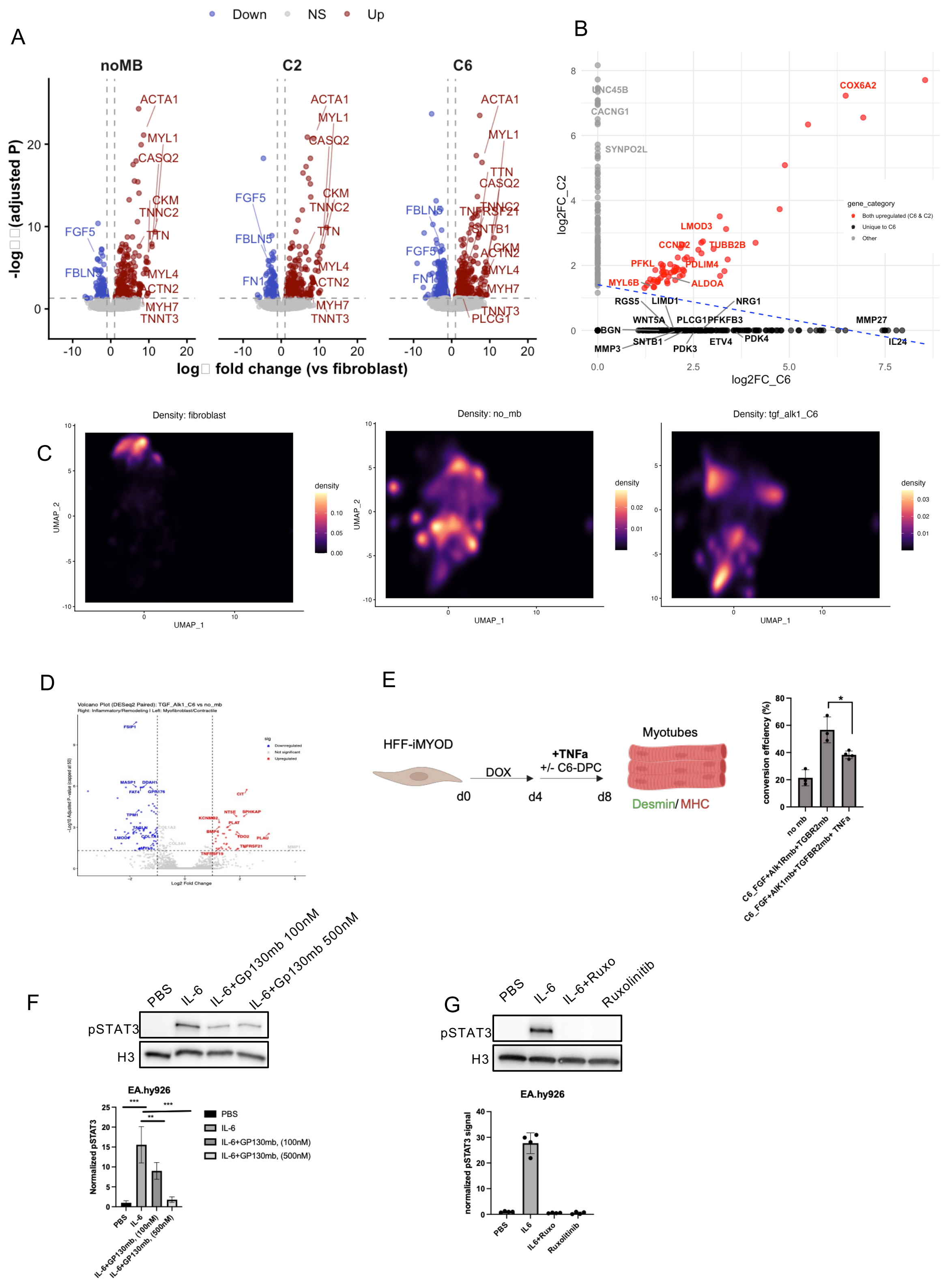

A
Down
NS
Up
noMB
C2
C6
ACTA1
MYL1
CASQ2
CKM
TNNC2
TTN
MYL4
ACTN2
MYH7
TNNT3
FGF5
FBLN5
FN1
TNFRSF21
SNTB1
PLCG1
log2FC_C2
log2FC_C6
B
UNC45B
CACNG1
SYNPO2L
COX6A2
LMOD3
CCND2
TUBB2B
PFKL
PDLIM4
MYL6B
ALDOA
RGS5
LIMD1
NRG1
WNT5A
PLCG1
PFKFB3
MMP27
BGN
SNTB1
ETV4
PDK4
IL24
MMP3
PDK3
gene_category
Both upregulated (C6 & C2)
Unique to C6
Other
C
Density: fibroblast
Density: no_mb
Density: tgf_alk1_C6
density
UMAP_1
UMAP_2
D
E
HFF-iMYOD
DOX
+TNFa
+/- C6-DPC
d0
d4
d8
Myotubes
Desmin/ MHC
conversion efficiency (%)
no mb
C6_FGF+Alk1Rmb+TGBR2mb
C6_FGF+Alk1mb+TGFBR2mb+ TNFa
F
PBS
IL-6
IL-6+Gp130mb 100nM
IL-6+Gp130mb 500nM
pSTAT3
H3
EA.hy926
Normalized pSTAT3
IL-6+GP130mb, (100nM)
IL-6+GP130mb, (500nM)
G
PBS
IL-6
IL-6+Ruxo
Ruxolinitib
pSTAT3
H3
EA.hy926
normalized pSTAT3 signal
IL6
IL6+Ruxo
Ruxolitinib

### Figure-EV4

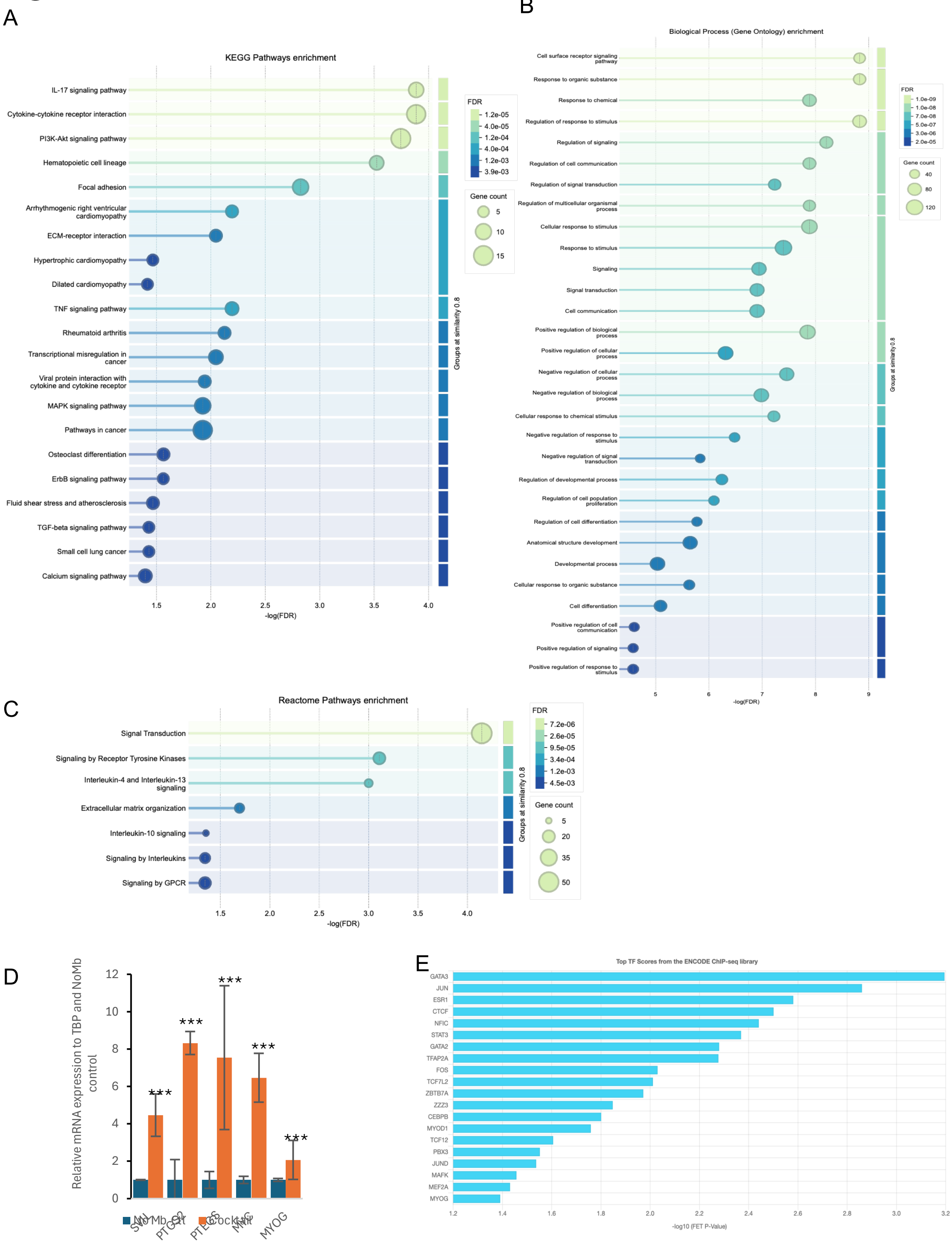

A
KEGG Pathways enrichment
IL-17 signaling pathway
Cytokine-cytokine receptor interaction
PI3K-Akt signaling pathway
Hematopoietic cell lineage
Focal adhesion
Arrhythmogenic right ventricular cardiomyopathy
ECM-receptor interaction
Hypertrophic cardiomyopathy
Dilated cardiomyopathy
TNF signaling pathway
Rheumatoid arthritis
Transcriptional misregulation in cancer
Viral protein interaction with cytokine and cytokine receptor
MAPK signaling pathway
Pathways in cancer
Osteoclast differentiation
ErbB signaling pathway
Fluid shear stress and atherosclerosis
TGF-beta signaling pathway
Small cell lung cancer
Calcium signaling pathway
1.5
2.0
2.5
3.0
3.5
4.0
-log(FDR)
Groups at similarity 0.8
FDR
1.2e-05
4.0e-05
1.2e-04
4.0e-04
1.2e-03
3.9e-03
Gene count
5
10
15
B
Biological Process (Gene Ontology) enrichment
Cell surface receptor signaling pathway
Response to organic substance
Response to chemical
Regulation of response to stimulus
Regulation of signaling
Regulation of cell communication
Regulation of signal transduction
Regulation of multicellular organismal process
Cellular response to stimulus
Response to stimulus
Signaling
Signal transduction
Cell communication
Positive regulation of biological process
Positive regulation of cellular process
Negative regulation of cellular process
Negative regulation of biological process
Cellular response to chemical stimulus
Negative regulation of response to stimulus
Negative regulation of signal transduction
Regulation of developmental process
Regulation of cell population proliferation
Regulation of cell differentiation
Anatomical structure development
Developmental process
Cellular response to organic substance
Cell differentiation
Positive regulation of cell communication
Positive regulation of signaling
Positive regulation of response to stimulus
5
6
7
8
9
-log(FDR)
Groups at similarity 0.8
FDR
1.0e-09
1.0e-08
7.0e-08
5.0e-07
3.0e-06
2.0e-05
Gene count
40
80
120
C
Reactome Pathways enrichment
Signal Transduction
Signaling by Receptor Tyrosine Kinases
Interleukin-4 and Interleukin-13 signaling
Extracellular matrix organization
Interleukin-10 signaling
Signaling by Interleukins
Signaling by GPCR
1.5
2.0
2.5
3.0
3.5
4.0
-log(FDR)
Groups at similarity 0.8
FDR
7.2e-06
2.6e-05
9.5e-05
3.4e-04
1.2e-03
4.5e-03
Gene count
5
20
35
50
D
Relative mRNA expression to TBP and NoMb control
12
10
8
6
4
2
0
***
***
***
***
***
SMN
PTGS2
PTGS
MMP
MYOG
NoMb-Ctrl
Clock-Mut
E
Top TF Scores from the ENCODE ChIP-seq library
GATA3
JUN
ESR1
CTCF
NFIC
STAT3
GATA2
TFAP2A
FOS
TCF7L2
ZBTB7A
ZZZ3
CEBPB
MYOD1
TCF12
PBX3
JUND
MAFK
MEF2A
MYOG
1.2
1.4
1.6
1.8
2.0
2.2
2.4
2.6
2.8
3.0
3.2
-log10 (FET P-Value)
